# Aminobisphosphonates reactivate the latent reservoir in people living with HIV-1

**DOI:** 10.1101/2023.02.07.527421

**Authors:** Marta Sanz, Ann Marie K. Weideman, Adam R. Ward, Matthew L. Clohosey, Susana Garcia-Recio, Sara R. Selitsky, Brendan T. Mann, Marie Anne Iannone, Chloe P. Whitworth, Alisha Chitrakar, Carolina Garrido, Jennifer Kirchherr, Alisha R. Coffey, Yi-Hsuan Tsai, Shahryar Samir, Yinyan Xu, Dennis Copertino, Alberto Bosque, Brad R. Jones, Joel S. Parker, Michael G. Hudgens, Nilu Goonetilleke, Natalia Soriano-Sarabia

**Affiliations:** Department of Microbiology Immunology and Tropical Medicine, the George Washington University, Washington DC, USA; Department of Biostatistics, University of North Carolina at Chapel Hill, North Carolina, USA; Department of Infectious Diseases, Weill Cornell Medicine, New York, USA; UNC HIV-1 Cure Center, University of North Carolina at Chapel Hill, North Carolina, USA; Lineberger Comprehensive Cancer Center, University of North Carolina at Chapel Hill, North Carolina, USA; Department of Genetics, University of North Carolina at Chapel Hill, North Carolina, USA; Microbiology & Immunology, University of North Carolina at Chapel Hill, North Carolina, USA

**Keywords:** HIV cure, latency reversing agents, HIV reservoir, shock and kill, γδ T cells, Vδ2 T cells, CD8+ T cells, IPDA, CyTOF.

## Abstract

Antiretroviral therapy (ART) is not curative due to the existence of cellular reservoirs of latent HIV-1 that persist during therapy. Current research efforts to cure HIV-1 infection include “shock and kill” strategies to disrupt latency using small molecules or latency-reversing agents (LRAs) to induce expression of HIV-1 enabling cytotoxic immune cells to eliminate infected cells. The modest success of current LRAs urges the field to identify novel drugs with increased clinical efficacy. Aminobisphosphonates (N-BPs) that include pamidronate, zoledronate, or alendronate, are the first-line treatment of bone-related diseases including osteoporosis and bone malignancies. Here, we show the use of N-BPs as a novel class of LRA: we found in *ex vivo* assays using primary cells from ART-suppressed people living with HIV-1 that N-BPs induce HIV-1 from latency to levels that are comparable to the T cell activator phytohemagglutinin (PHA). RNA sequencing and mechanistic data suggested that reactivation may occur through activation of the activator protein 1 signaling pathway. Stored samples from a prior clinical trial aimed at analyzing the effect of alendronate on bone mineral density, provided further evidence of alendronate-mediated latency reversal and activation of immune effector cells. Decay of the reservoir measured by IPDA was however not detected. Our results demonstrate the novel use of N-BPs to reverse HIV-1 latency while inducing immune effector functions. This preliminary evidence merits further investigation in a controlled clinical setting possibly in combination with therapeutic vaccination.

## INTRODUCTION

Antiretroviral therapy (ART) is not curative due to the existence of cellular reservoirs of latent HIV-1 that persist during therapy (1). Current research efforts to cure HIV-1 infection include “shock and kill” strategies to disrupt latency using small molecules or latency-reversing agents (LRAs) to induce expression of HIV-1 enabling cytotoxic immune cells to eliminate infected cells (2). Despite the appealing nature of the theory behind the “shock and kill” strategy for HIV-1 cure, the latency reversing agents (LRAs) that have progressed to clinical trials to date have not achieved a sustained decrease of the viral reservoir despite demonstrating *in vivo* induction of HIV-1 RNA transcription or transient detectable plasma viremia (3, 4). Both the limited availability of Federal Drug Agency (FDA)-approved LRA compounds and their modest success to date urges the HIV-1 cure field to identify additional drugs for pre-clinical and clinical testing.

Aminobisphosphonates (N-BPs), like pamidronate (PAM), zoledronate (Zol), or alendronate (ALN), function by inhibiting the enzyme farnesyl pyrophosphate synthase in the mevalonate pathway, which results in 1) inhibition of the production of the key intermediate farnesyl pyrophosphate (FPP) and 2) accumulation of upstream isopentenyl pyrophosphate(5, 6). FPP is essential for protein prenylation, a post-translational modification that consists of the transfer of FPP groups to proteins involved in cell signaling, including GTPases from the Ras family (7, 8). Interestingly, statins that inhibit upstream of IPP production have been shown to inhibit HIV-1 viral replication (9–11). Upstream accumulation of IPP also leads to the exclusive activation of the more frequent subpopulation of circulating gamma delta (γδ) T cells, Vδ2 T cells, that specifically recognize IPP in a T cell receptor (TCR)-dependent manner (12–14). Vδ2 T cells are potent cytotoxic effectors against virally infected or malignant cells (15–17), and we previously reported the capacity of *ex vivo* expanded Vδ2 T cells to target and eliminate latently HIV-1-infected resting CD4 T cells upon *ex vivo* latency reversal (18, 19). In addition, activated Vδ2 T cells have a critical adjuvant activity by inducing antigen-experienced and naïve CD8 T cell responses (20–24). We hypothesized that N-BPs exert a dual effect by inducing both HIV-1 reactivation and concomitant immune responses through activation of Vδ2 T cell cytotoxic and adjuvant effects that could lead to elimination of HIV-1 reservoirs.

Here, we show that *ex vivo* exposure to N-BPs reverses HIV-1 latency most likely through the activator protein 1 (AP-1) signaling pathway. *In vivo* treatment of ART-suppressed people living with HIV (PLWH) with ALN induced viral reactivation although a significant reduction in the reservoir size was not detected in our small cohort. This study provides evidence of the novel use of N-BPs as LRAs and warrants further investigations with the specific objective to assess latency reversal and infected cell clearance in a larger cohort.

## RESULTS

### N-BPs induce *ex vivo* HIV-1 reactivation

#### Ex vivo study population

N-BPs function by inhibiting FDPS enzyme leading to activation of Vδ2 T cells, although potency may vary (25–27). PAM was primarily used for our *ex vivo* studies and compared to Zol and ALN when sufficient cells were available. We utilized N-BP concentrations (PAM at 2.5μg/mL and 25μg/mL, Zol at 1μg/mL, and ALN at 2.5μM) that mimic plasma concentrations achieved in the blood with FDA-approved doses (28, 29), and analyzed HIV-1 cell-associated RNA (caRNA) and replication competent HIV-1 production by quantitative viral outgrowth assay (QVOA) after six hours of N-BP exposure. In some experiments, cells were treated in parallel with VOR 350nM (for HIV-1 caRNA) or 500nM (for QVOA).

#### HIV-1 cell-associated RNA

The *ex vivo* capacity of N-BPs to reverse HIV-1 latency was analyzed in 24 to 72 million resting CD4 (rCD4) T cells from 23 PLWH on suppressive ART (Supplementary Table 1). Untreated biological replicates of 1×10^6^ cells, or treated with N-BPs, or with PHA and IL-2 for 6 hours were compared. Spontaneous production of HIV-1 without treatment was measurable in almost all individuals despite long term suppression (Supplementary Fig. 1), and was inversely correlated with the CD4 T cell nadir (Figure 1A). Fold change viral transcription induction, mediated by PAM (14 donors at 2.5μg/mL), Zol (9 donors) and ALN (5 donors), was comparable to the positive control PHA+IL-2 (Figure 1B). A higher 25µg/mL PAM concentration was also tested in 11 donors, and was compared to vorinostat (VOR) in seven donors. PAM at 25μg/mL induced slightly higher reactivation than 500nM VOR (Supplementary Fig. 1C and 1D).

**Figure 1:**
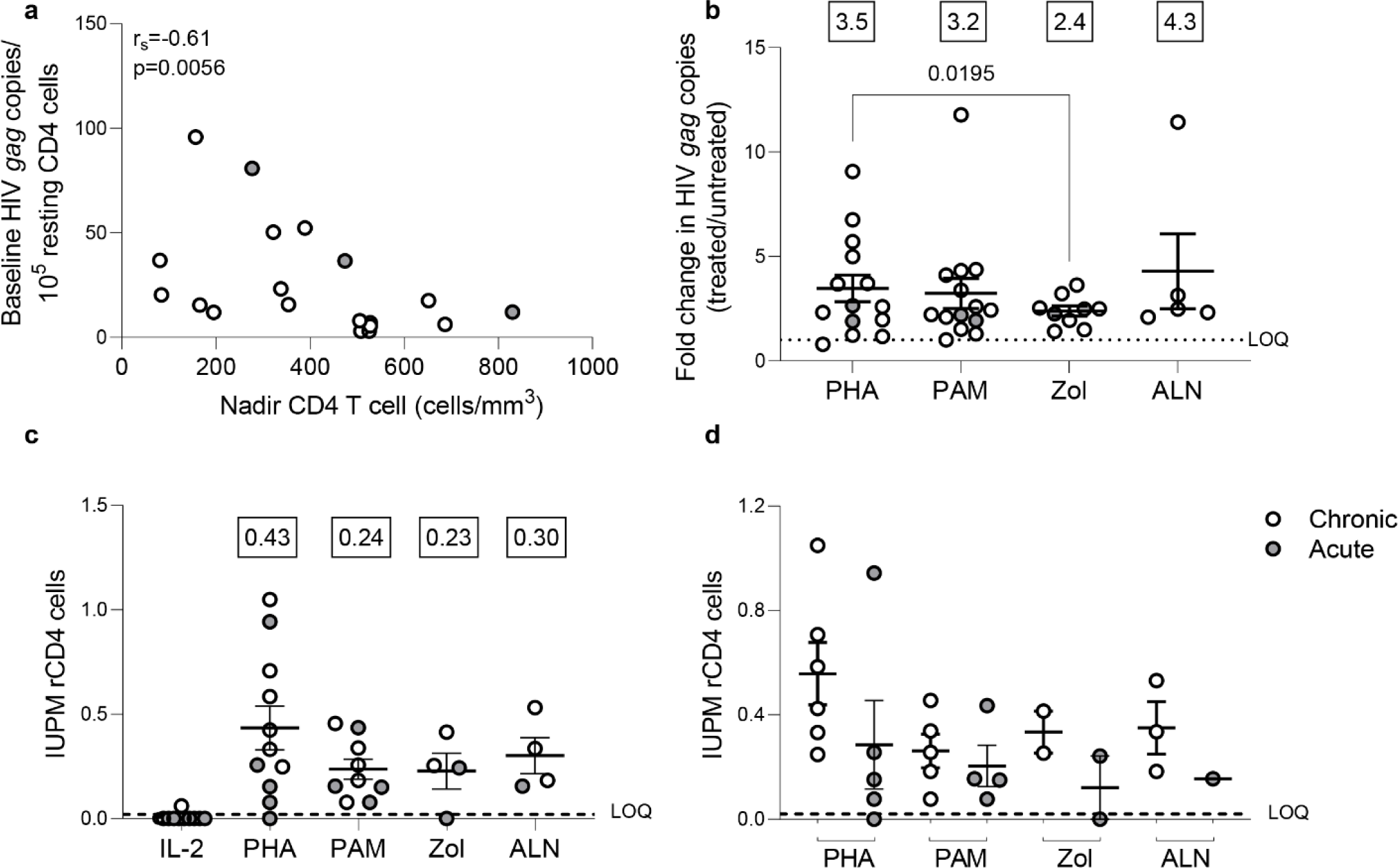
N-BPs induce reactivation of latent HIV *ex vivo*. **(A)** Inverse correlation between spontaneous HIV production and CD4 T cell nadir in 19 individuals with available CD4 T cell nadir data (Spearman’s rank correlation). **(B)** Fold change induction of HIV caRNA levels. Comparable capacity of PAM (N=14), Zol (N=9), and ALN (N=5) to the positive control PHA + IL-2 (N=14) to induce production of HIV *gag* copies from isolated rCD4 T cells (p>0.05, Wilcoxon signed-rank test). Each symbol represents one individual for whom a mean of 6-15 replicates of 1×10^6^ million cells per condition were assayed. Boxed values in B and C are the mean fold change for each condition. Infectious Units per Million (IUPM) rCD4 T cells from QVOA **(C)** in all donors and **(D)** according to treatment initiation in the acute or chronic phase of HIV-1 infection. Cells were treated with PAM, ALN and Zol and compared to PHA (PHA+IL-2). IL-2 alone was used as a control of the assay. (Only p-values < 0.05, generated by a Wilcoxon signed-rank test, are displayed). Boxed values are the mean IUPM for each condition. Mean± standard error of the mean (SEM) is represented in (B) and (C). LOQ, limit of quantification. Filled grey circles represent donors treated during the chronic phase of infection and opened circles are donors treated during the acute phase of infection.

#### Replication-competent HIV-1

To confirm that N-BPs induced production of non-defective, replication-competent HIV-1, rCD4 T cells from PLWH on ART were used in QVOAs using a minimum of 13 million rCD4 T cells in each treatment condition. PHA+IL-2 induced production of replication competent HIV-1 in 10 of 11 individuals with a group mean of 0.43 IUPM (range: 1.5×10^-8^ to 1.05), PAM produced 0.24 IUPM (range: 0.08 to 0.46), Zol 0.23 IUPM (range: 1.5×10^-8^ to 0.41), and ALN 0.30 IUPM (range: 0.16 to 0.53) (Figure 1C). The effects of PAM, ALN and Zol were comparable to PHA (Figure 1C). A trend towards a lower IUPM in rCD4 T cells from donors treated during the acute phase of HIV infection was observed (Figure 1D). Our results demonstrate a potent capacity of N-BPs to reverse HIV-1 latency *ex vivo*.

## Mechanism of action of N-BPs on viral reactivation

To explore pathways involved in N-BPs reversing HIV-1 latency, we performed RNA-seq on *ex vivo* untreated or 2.5μg/mL PAM-treated paired samples from six ART-suppressed PLWH. To exclude changes in gene expression, due to toxicity, a dose-curve analysis of PAM toxicity was performed showing that the selected dose did not impact cell viability (Supplementary Fig. 2). Next, to confirm the functional effect of N-BPs at selected concentrations in our experiment, we analyzed whether protein prenylation of Ras was reduced upon N-BP exposure. Our results showed that treatment with PAM, Zol and ALN decreased the expression of active Ras (GTP-Ras) after overnight incubation by 3.1-fold, 4.7-fold and 6.6-fold compared to media only control, respectively, indicating inhibition of protein prenylation (Figure 2A). RNA-seq analysis identified 1,799 genes differentially expressed between samples treated with PAM and untreated controls (Figure 2B, Supplementary Table 2). We next tested for differentially expressed gene modules using Gene Set Variation Analysis and linear regression (GSVA, Supplementary Table 3) (30). Consistent with the effect of N-BPs decreasing the production of cholesterol (31), the cholesterol synthesis pathway (GO_Regulation of cholesterol biosynthetic pathway) was decreased in the samples treated with PAM compared to media only control (Supplementary Fig. 3), confirming the specificity of PAM and validating the RNA-seq data. The top differentially expressed pathway was the response to bacterial lipoprotein, and identified TLR2 as one of the genes upregulated upon PAM treatment (p= 4.6×10^-15^; Figure 2C and Supplementary Table 3). To confirm the effect of PAM on this pathway, we performed time-course surface TLR2 expression studies by flow cytometry in isolated CD4 T cells. In accordance with our RNA-seq data (Supplementary Table 2), we detected an early upregulation of surface TLR2 expression after four and six hours of incubation, followed by a downregulation after overnight exposure to N-BPs (Figure 2D). Next, we used the Predicting Associated Transcription factors from annotated affinities (PASTAA)(32) software to predict transcription factors (TF) associated with PAM-mediated transcriptional changes. The top-8 TFs associated with the PAM-mediated transcriptional changes were NKX2-1, IRF-8, ETV4, POU2F1, two subunits of AP-1 (FOSB and JUN), ELF-4 and Cart-1. STRING pathway analysis of interactions of these TFs with the GTPases activated upon inhibition of protein prenylation (Rac, Cdc42 and Rho) revealed a unique cluster of these GTPases with JUN (c-Jun) (Figure 2E), that has been previously shown to be involved in viral reactivation from latency (33, 34). Since c-Jun transcriptional activation depends on phosphorylation at Ser73 of the transactivation domain for transcriptional activity (35) we next analyzed the phosphorylation status of c-Jun at this residue upon 15 minutes of exposure to N-BPs and PMA as a positive control. Phosphoflow analysis showed consistent induction of c-Jun phosphorylation mediated by Zol, and to a lesser degree by PAM and ALN (Figure 2F). This increase in phosphorylation by N-BPs was not observed in p65, the active subunit of NFκB (Figure 2G).

**Figure 2:**
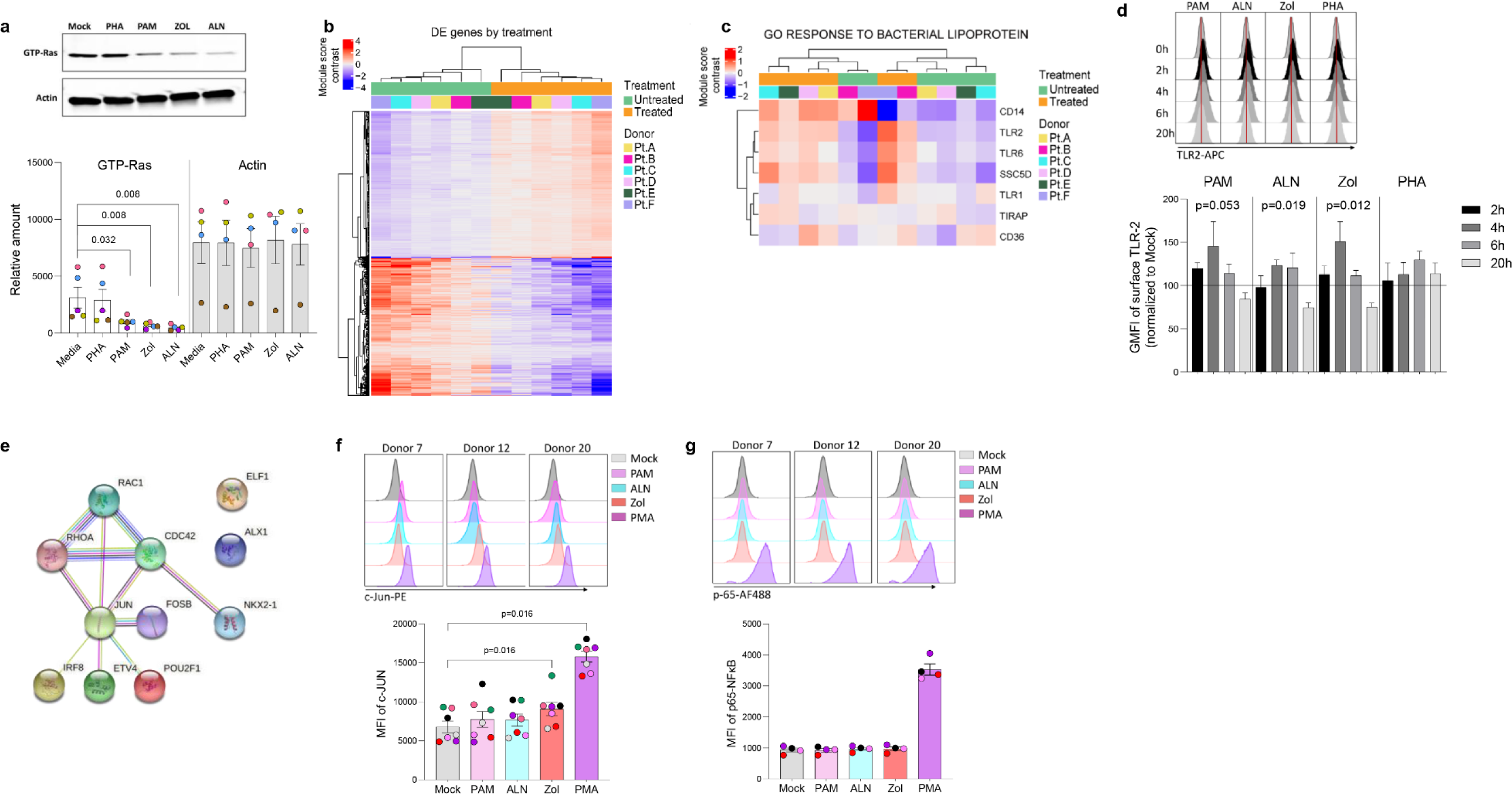
N-BP’s mechanism of action. *Ex vivo* isolated rCD4 T cells from ART-suppressed PLWH were treated with 2.5μg/mL PAM or left untreated for 6 hours, washed, and RNA extracted for RNA-seq analysis. **A)** Representative western blotting of activated Ras (GTP-Ras) and Actin in isolated CD4+ T cells and integrated band density analysis (ImageJ) in 5 independent samples after overnight incubation with PHA, PAM, ALN or Zol (p-values generated by a Mann-Whiney U test). **B)** Heatmap displaying the 1,799 differentially expressed (DE) genes (Wald statistic conditioned on donor, q-value < 0.05, and base mean >10) comparing *ex vivo* untreated samples versus PAM-treated samples; color represents relative expression. Columns and rows were organized by unsupervised hierarchical clustering. Pt, participants. **C)** Heatmap displaying genes in the module “GO Response to bacterial lipoprotein”. Columns and rows are organized by unsupervised hierarchical clustering. **D)** Modulation of TLR2 expression on *ex vivo* isolated CD4+ T cells upon treatment with ALN, PAM, Zol or PHA. Overlaid histograms from one representative donor showing mean fluorescence intensity (GMFI), and normalized data to the untreated control at each time point from four different donors (Friedman test). **E)** STRING pathway analysis. Representative histograms from three different donors showing MFI expression of **F)** c-Jun phosphorylation at Ser73 and mean expression of c-Jun phosphorylation (N=7) and **G)** p-65 NFκB subunit phosphorylation at Ser529, and mean expression of p-65 (N=4), upon exposure to N-BPs or PMA.

## Effect of ALN on the HIV-1 reservoir

### In vivo study population

To examine the effects of N-BPs *in vivo*, we obtained retrospective PBMCs and plasma samples (baseline, and weeks 2, 24 and 48 post-intervention) from the clinical trial ACTG A5163 that previously assessed the efficacy of ALN vs. placebo to increase bone mineral density in PLWH on ART (36). Eligibility criteria for the ACTG A5163 trial included two consecutive measures of viral suppression (≤5,000 RNA copies by any quantitative HIV viral load assay) prior to enrollment. Our criteria to obtain banked samples were viral suppression and availability of stored samples with more than 5 million PBMCs at study entry, and at least one follow-up sample collection. Plasma samples were used to confirm viral suppression at study entry and at follow-up time points. We received samples from 57 participant and thawed PBMCs demonstrated acceptable viability, integrity, cell numbers and viral suppression (< 40copies/mL) in 44 of the 57 samples, 23 from the ALN group and 21 from the placebo group (Supplementary Table 4). Samples were randomly assigned to perform the intact proviral DNA assay (IPDA) in nine participants from the ALN group and seven from the placebo group (Supplementary Table 5). Quantification of HIV-1 caRNA, mass cytometry, and HIV-specific T cell responses by ELISPOT were performed in 15 ALN-treated participants and 14 from the placebo group (Supplementary Table 6). All experiments were assayed blinded.

### ALN treatment reactivates the latent HIV-1 reservoir in PLWH

Unspliced HIV-1 *gag* caRNA levels were measured in 14 participants, nine individuals treated with ALN and five who received placebo. In each individual, between six to 15 replicates of 1×10^6^ PBMCs were assayed. Baseline levels were comparable between the two groups (Figure 3A) and were not associated with baseline CD4 or CD8 T cell counts, time on ART or age (p>0.05 for all comparisons). Seven participants (five treated with ALN) were women and had lower baseline caRNA levels than men (Figure 3B). Mean longitudinal caRNA levels remained comparable between the two interventions (Figure 3C) and were lower in women compared to men (Figure 3D). However, the lack of difference between ALN and placebo groups was due to variable HIV-1 caRNA dynamics in the ALN group, as observed when analyzing individual donors. In eight of nine participants, we observed differential effects of ALN treatment on HIV-1 caRNA (Figure 3E). caRNA levels in ALN-treated participants 101, 102 and 103 showed a positive trend in caRNA levels from basal to available follow up time points, while participant 104 had a transient increase at week 2 after the intervention. By contrast, in ALN-treated participants 105, 106, 107 and 108, HIV-1 caRNA expression decayed from baseline compared to later time-points following ALN treatment (Figure 3E). HIV-1 caRNA trends remained stable within the five donors analyzed from the placebo group (Figure 3F). In three participants (101 with a positive trend, and 105 and 107 with a negative trend), total HIV-1 DNA was quantified (Figure 3G). All three exhibited a sustained decay of total HIV-DNA. Participant 107 also exhibited a decay in intact provirus and defective DNA species (Supplementary Fig. 4). Given the complex dynamics of HIV-1 caRNA (37–39), we may be capturing initial latency reversal after which there may be clearance and decay, therefore results may be reflecting different timing and duration of the measurement.

**Figure 3:**
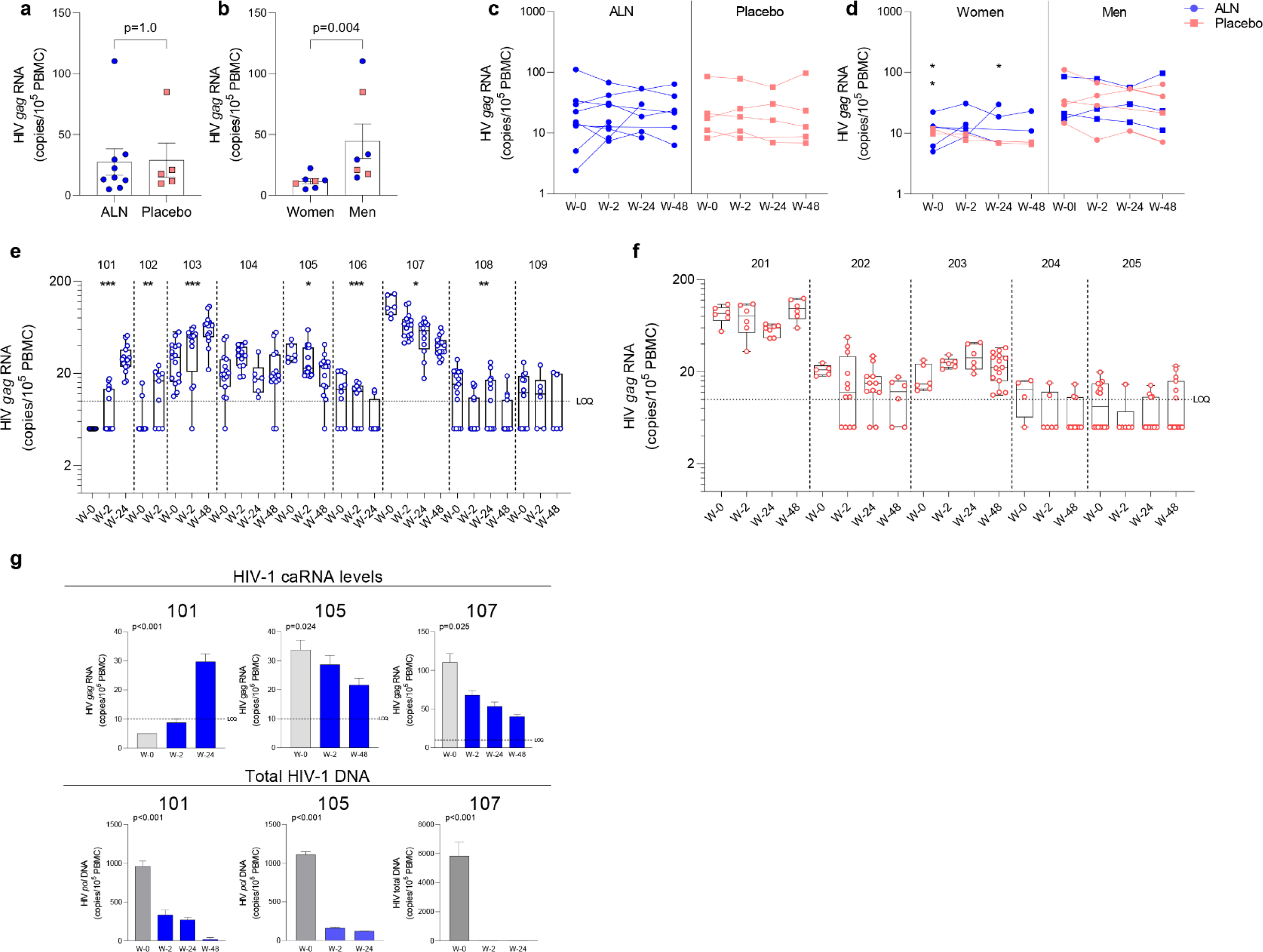
Effect of *in vivo* ALN treatment on unspliced HIV *gag* caRNA. Baseline caRNA levels according to **A)** treatment, ALN or placebo, and **B)** biological sex, woman or man. (Mann-Whitney U test). The ALN group is represented in blue, and the placebo group is represented in pink. Longitudinal HIV caRNA levels according to **C)** treatment, ALN or placebo (Holm-Bonferroni-adjusted p=1.0, at baseline, week 2, and week 24 for ALN and placebo; Mann-Whitney U test), and **D)** biological sex, women and men (Holm-Bonferroni-adjusted p=0.01 at baseline, p=0.07 at week 2, p=0.07 at week 24. N=2 at week 48, precluding statistical analysis; Mann-Whitney U test). (N=3 and N=4 for ALN vs placebo; N=2 and N=5 for Women vs Men). Individual HIV caRNA copies/10^5^ PBMCs in participants who took **E)** ALN or **F)** placebo. Box and whiskers plots represent the min to max values. 5-15 biological replicates of 1×10^6^ cells were performed per individual. Mann-Kendall trend test, *p<0.05, **p<0.01, ***p<0.001. **F)** Paired caRNA levels and total HIV DNA levels was available in participants 101 and 105 (*pol* HIV DNA by ddPCR) and 107 (by IPDA). HIV DNA copies were “0” at weeks 2 and 24. Baseline values are presented in grey and post-intervention weeks in blue. P-values are calculated using a Mann-Kendall trend test.

### Effect of ALN on the viral reservoir in PLWH

The intact proviral DNA assay (IPDA) was performed on PBMCs, and no assay detection failures were observed with all 15 participants showing amplification of the Packaging Signal (Ψ) and *env* regions. Both ALN and placebo groups had comparable baseline mean HIV-1 DNA levels as follows: intact provirus (356 vs. 471, p=0.76), total HIV-1 DNA (3,560 vs. 3,483, p=1.0), total defective HIV-1 DNA (the sum of hypermutated, 3’defective, and 5’ defective -3,204 vs. 3,012, p=0.92), hypermutated / 3’ defective (1828 vs. 1607, p=1.0), and 5’ defective (1377 vs. 1405, p=1.0). The mean proportion of intact proviruses out of total HIV-1 DNA at baseline was 11.5% for all participants, 9.9% for the ALN group and 13.5% for the placebo group (Figure 4A). Intact proviral and total HIV-1 DNA levels correlated at baseline (Figure 4B) and at subsequent time points after the intervention (not shown).

**Figure 4:**
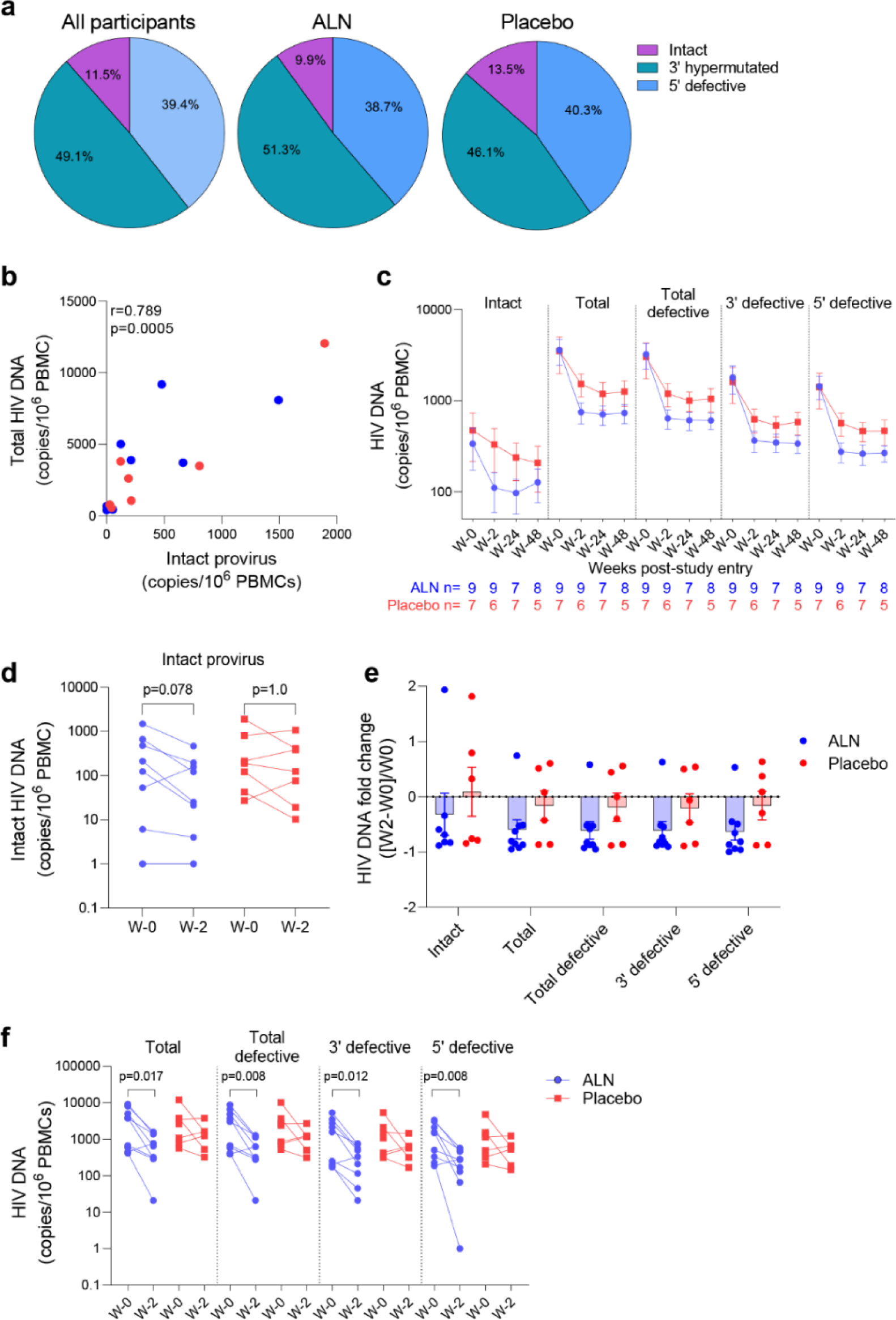
Effect of ALN on the viral reservoir measured by IPDA. **A)** Proportion of defective HIV DNA and intact proviruses. Numbers represent proportion of total. **B)** Correlation between intact proviral and total HIV-1 DNA levels at study entry for ALN (blue) and placebo (red) participants (Spearman’s correlation). **C)** Longitudinal HIV DNA levels in participants from the ALN and placebo groups. Mean ± standard error of the mean (SEM) is presented. p>0.05 for all comparisons for both ALN and placebo, Mann-kendall test. **D)** Comparison of intact proviral levels between baseline and week 2 after intervention in the ALN and placebo groups, Wilcoxon signed-rank test. **E)** HIV-1 DNA fold change at week 2 with respect to week 0 calculated as ((week-2–week-0)/week-0) is presented, p>0.05 for all comparisons, Mann Whitney U-test. **F)** Changes in total, total defective, hypermutated / 3’ defective and 5’ defective HIV-1 DNA from baseline to week 2 after intervention in ALN and placebo groups, Wilcoxon signed-rank test.

HIV-1 DNA copies decayed from baseline to week 2 and remained stable in longitudinal measures until week 48 in both groups (Figure 4C and supplementary table 7). Changes in total and defective HIV-1 DNA levels were not associated with baseline CD4 or CD8 T cell counts, time on ART or age (Supplementary Table 8). A trend towards a decay from baseline to week two in intact provirus was observed in the ALN group which was not observed in the placebo group (p=0.078 vs. p=1.0, Figure 4D). However, as three individuals from the placebo group had a similar proportional decay in intact proviral DNA levels compared to the three participants with largest decays in the ALN group, a larger cohort is required to confirm this preliminary trend. Similarly, fold changes from baseline to week two after intervention were similar between ALN and placebo groups for all HIV DNA species (Figure 4E). Finally, we observed differences in changes from baseline to week two in total and defective HIV-1 DNA only in the ALN group (Figure 4F). Comparable HIV-1 DNA copies over time in both groups (Figure 4C), the small and uneven sample size (n=9 for the ALN group vs. n=7 for the placebo group), and the variability of the assay (Supplementary Fig. 5) precludes drawing conclusions regarding the effect of ALN on the viral reservoir. These results warrant further investigation to analyze whether N-BP-mediated viral reactivation culminates in a reduction of the viral reservoir.

## Effect of ALN on immune cells

We performed mass cytometry in eight participants from the ALN group and six from the placebo group. Over the course of ALN treatment frequencies of Vδ2 and Vδ1 T cells remained stable whereas effector memory (EM)-CD8 T cell frequencies decreased. (Figure 5A). All the other cell populations analyzed showed comparable baseline and longitudinal frequencies (Supplementary Figs. 6 and 7). Higher granzyme B (GzmB) production from Vδ2 T cells, Vδ1 T cells and CD8 T cells compared to the placebo group after 24 weeks of intervention were detected (Figure 5B). Comparison by biological sex revealed higher baseline GzmB production from Vδ2 T cells, Vδ1 T cells, and CD8 T cells in women than in men, although men experienced a longitudinal increase that reached comparable levels to women (Figure 5C). Although we did not detect an effect of ALN on HIV-1 specific T cell responses (Supplementary Fig. 8), there was evidence for changes in Vδ2 T cell functionality (Figure 5B) that led to an induction of CD8 T cell functions including decreased peripheral EM-CD8 T cells (Figure 5A) and increased apoptotic GzmB granule production (Figure 5B).

**Figure 5:**
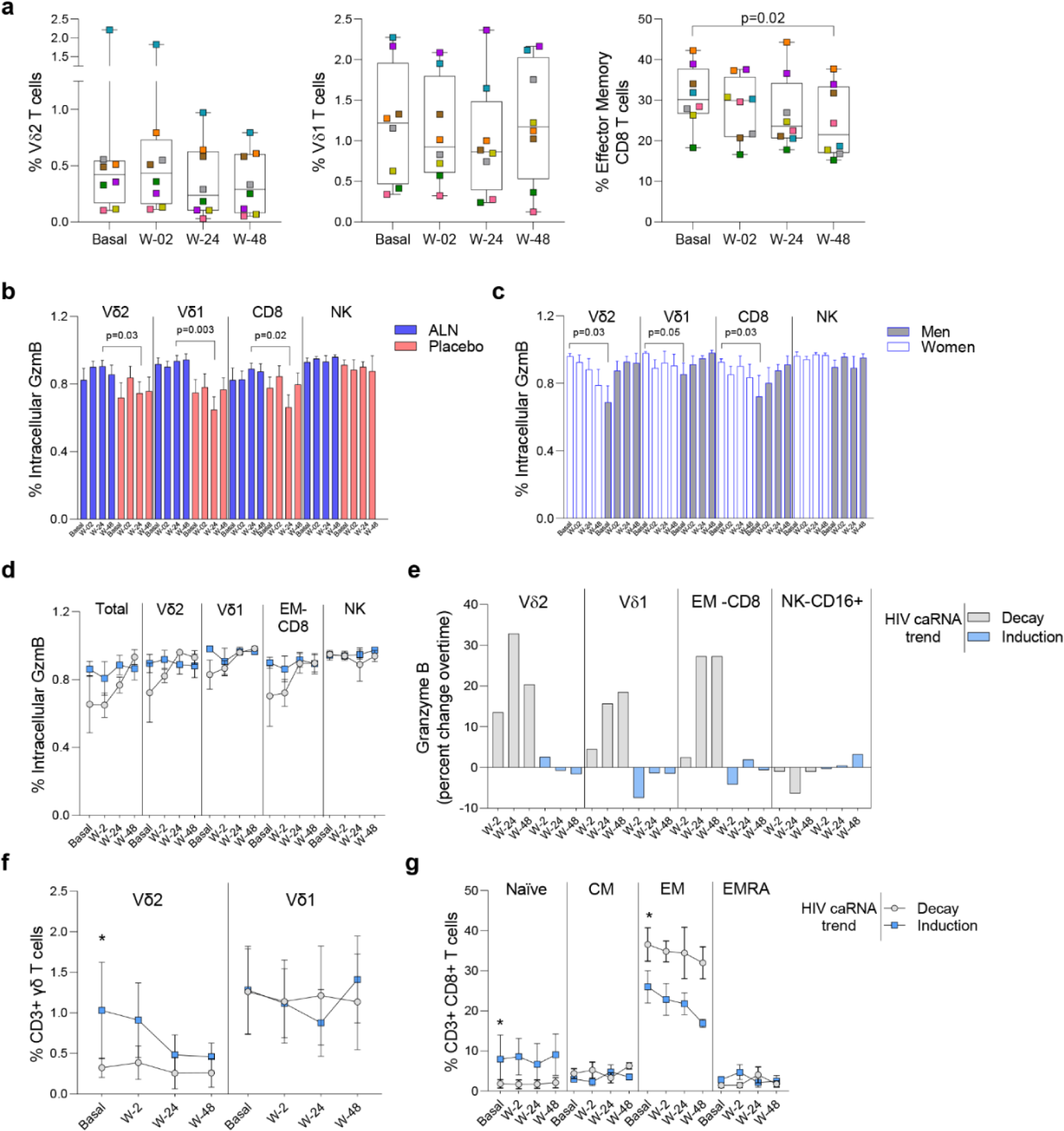
Impact of ALN treatment on circulating immune cells’ phenotype and functional markers. **A)** Frequency of circulating Vδ2 T cells, Vδ1 T cells and Effector Memory CD8 T cells. Boxplots display first quartile, median, and third quartile with whiskers range from the minimum to maximum values. P-values are from a Wilcoxon-signed rank test and adjusted for multiple comparisons using Holm-Bonferroni. Intracellular production of Granzyme (Gzm) B according to **B)** the intervention and **C)** biological sex. Mean ±SEM in eight participants is represented**. D)** Total and effector cell population’s intracellular production of GzmB according to the HIV caRNA trend (grey, participants in whom HIV-1 caRNA decay was observed (N=3), and blue, participants in whom HIV-1 caRNA induction was observed (N=3). **E)** Percent change over time in GzmB, IFN-γ and TNF-α. Values below zero represent reduction from baseline and values above zero represent increase from baseline. Circulating frequencies of **F)** γδ T cell populations **G)** CD8 T cell subpopulations. Mean ± SEM is presented. Mann-Whitney U test. Mass cytometry data on 8 participants treated with ALN and 7 from the placebo group. Mass cytometry data represents the frequency of alive events measured.

### Increased Granzyme B production associates with caRNA decay

To understand the dynamics of HIV-1 caRNA observed in the ALN group (Figure 3E) we compared GzmB, IFN-γ and TNF-α production from three individuals with an induction of HIV-1 caRNA copies, and three with a decay in HIV-1 caRNA levels. Most noticeable changes included a consistent longitudinal increase in GzmB production (Figure 5D), and calculated percent changes over time showed that Vδ2 T cells, Vδ1 T cells, and CD8 T cells contributed to increasing GzmB production (Figure 5E). In addition, in participants with a decay in HIV-1 caRNA levels we observed constant Vδ2 T cell frequencies over time, despite being lower than in participants with increased HIV-1 caRNA levels (Figure 5F), lower frequency of naïve CD8 T cells and higher frequency of EM-CD8 T cells (Figure 5G). Other comparisons are shown in Supplementary Fig. 9. These immune differences may contribute to the variable HIV-1 caRNA slopes and warrants further investigation in a larger cohort.

## MATERIALS AND METHODS

### Donors

For *ex vivo* studies, frozen samples from PLWH on ART with undetectable plasma viral load (<50 copies/mL) for at least one consecutive year before inclusion in this study were obtained at the University of North Carolina, USA. Participants’ characteristics have been previously reported (36). Frozen peripheral blood mononuclear cells (PBMCs) and plasma samples from the AIDS Clinical Trial Groups (ACTG) A5163 clinical trial performed in 2004 were requested for *in vivo* analyses (36). The only criteria to obtain the samples was that participants had to be virally suppressed and have cell availability of at least 5 million in the baseline and one follow-up timepoint. Briefly, the objective of that study was to assess the effect of once-weekly oral administration of ALN on bone mineral density. A5163 was a phase II, double blind, placebo-controlled study where chronic ART-suppressed HIV-1-seropositive participants were randomized into ALN and placebo interventions. Both arms took calcium carbonate and vitamin D and were followed for 48 weeks. PBMCs and plasma samples were obtained and stored prior to the intervention (basal) and at weeks 2, 24, and 48 post-intervention. All assays were performed blinded as per treatment status.

### Drugs and Compounds

PAM, Zol and ALN were used at concentrations that mimic plasma concentrations achieved in the blood with FDA-approved doses (28, 29). PAM was used at 2.5μg/mL or 25μg/mL, Zol at 1μg/mL, and ALN at (2.5µM). In some experiments, we treated cells in parallel with 350nM VOR (for HIV-1 caRNA levels quantification), or 500nM (for replication-competent HIV-1 determination). PAM at 2.5μg/mL and 25μg/mL was able to similarly induce HIV-1 caRNA and replication competent HIV-1-1, and was compared to VOR in some experiments.

### Quantitative Polymerase Chain Reaction (qPCR)

<u>For *ex vivo* analysis</u> of HIV-1 caRNA levels, magnetically isolated rCD4T cells from PLWH on suppressive ART were exposed to N-BPs, PHA and IL-2, or left untreated. In some experiments, cells were exposed VOR. After six hours of exposure, rCD4 T cells were washed and plated in 6-15 replicates of 1×10^6^ cells per condition, depending on cell availability. After further washes, cells were pelleted and stored at -80°C, until RNA isolation. <u>For the *in vivo* part of the study</u>, thawed PBMCs were rested overnight, plated in 3-5 replicates of 1×10^6^ cells (depending on cell availability), and stored at -80°C. Automated RNA extraction was performed (KingFisher, ThermoFisher Scientific), cDNA was synthesized using standardized protocols, and qPCR performed in triplicate using specific and validated primers, probes, and standard curves to amplify unspliced HIV-1 *gag*, as previously described (40). Results were expressed as HIV-1 *gag* copies per 10^5^ rCD4 T cells for the *ex vivo* analysis and 10^5^ PBMCs for the *in vivo* analysis. The limit of detection of the PCR was 10 HIV-1 *gag* RNA copies/10^5^ cells.

### Quantitative Viral Outgrowth Assay (QVOA)

HIV-1 p24 protein production (replication-competent virus as measure of infectious particles production) was analyzed in quantitative viral outgrowth assays (QVOA) (19, 41) after rCD4 T cells’ *ex vivo* exposure to N-BPs PAM and Zol. Briefly, magnetically isolated rCD4 (CD3+/CD4+/CD69-/ CD25-/ HLA-DR-) T cells (Stemcell Technologies) were rested in the presence of antiretroviral drugs (10nM Raltegravir and 20nM Efavirenz). Cells were washed, counted and plated in limiting dilution from 1×10^6^ to 0.1×10^6^ cells. Cells were left untreated or exposed 24 hours to PHA and IL-2, or to N-BPs PAM, Zol and ALN. After washing, isolated PHA-activated CD4 cells from uninfected individuals were added twice during the 19-day culture as targets of new infections to outgrowth the virus. After 15 days of culture, supernatants were harvested and HIV-1p24 ELISA performed and confirmed at day 19 of culture. The frequency of infection was calculated and expressed as infectious units per million (IUPM) cultured cells, as explained in Statistical Methods.

### RNA-Sequencing

In order to explore the mechanism of action of N-BPs related to HIV latency reactivation, RNA was isolated from rCD4 T cells exposed to PAM or VOR for 6 hours, or left untreated.

#### RNA isolation and quality control

RNA was isolated using the automated KingFisher (Thermofisher Scientific, Waltham, MA). All RNA samples were assayed for integrity, concentration, and fragment size on a TapeStation system (Agilent, Inc. Santa Clara, CA).

#### RNA-seq library preparation and sequencing

Library was constructed using Truseq RNA Exome library construction following the steps: Fragmenting RNA, Synthesizing 1st Strand cDNA, Synthesizing 2nd Strand cDNA, Adenylating 3’ Ends, Ligating Adaptors, Enriching DNA fragments, Hybridizing Probes to library, capturing Hybridized Probes, performing a 2nd Hybridization, performing a 2nd capture, Clean-up of captured library was performed via AMPure XP Beads followed by amplification of the enriched library, and a final clean-up of amplified enriched library using AMPureXP beads. Libraries were prepared on an Agilent Bravo Automated Liquid Handling System. Indexed libraries were prepared and run on HiSeq4000 to generated paired end 75 base pair reads which generated approximately 90 million reads per sample library with a target of greater than 90% mapped reads.

#### RNA-seq analysis

RNA-seq reads were aligned to the human genome GRCh38.d1.vd1 from Genomic Data Commons using STAR v2.4.2a (parameter: --quantMode TranscriptomeSAM, default mismatches and multi-maps) (42). Transcripts were quantified using Salomon v0.6 (PMID: 28263959, parameter: -l IU). DESeq2 was used for normalization and determination of differentially expressed genes (43). To determine differentially altered pathways, we computed a module score using Gene Set Variation Analysis (GSVA, PMID: 23323831) using the residuals of the gene expression after adjusting out the donor variation and linear regression for the GO modules from mSigDB, version 7 (30). Estimates and p-values were calculated for each gene module by fitting to a generalized linear mixed effects model.

### Phosphoflow cytometry

Phosphorylation status of p65 and c-JUN were analyzed by flow cytometry in isolated CD4 T cells (StemCell Technologies) as previously described (44). Briefly, cells were stimulated for 15 minutes at 37°C in the absence or presence of 2.5µM ALN, 2.5µg/mL PAM, 1µg/mL ZOL or 50nM PMA as a positive control. Cells were washed and stained with a viability dye (Zombie Aqua Fixable viability kit, BioLegend) and fixed with 100µl of pre-warmed Cytofix Fixation Buffer (BD Biosciences) for 10 minutes at 37°C. Cells were permeabilized while vortexing with 100 µL of prechilled Perm Buffer III (BD Biosciences) and incubated for 30 minutes on ice. After incubation, cells were washed and stained with anti-NFkB p65 (pS529, clone K10-895.12.50, BD Biosciences) or Phospho-c-Jun (Ser73, clone D47G9, Cell Signaling Technology) for 16 hours at 4°C. After washing, samples were acquired on a BD LSR Fortessa TM X-20 instrument (BD Biosciences) and analyzed using Flowjo (FlowJo v.10.8.1).

### Intact Proviral DNA Assay (IPDA)

Total DNA was extracted from frozen cell pellets of 0.3-1×10^6^ PBMCs (Qiagen, Germantown, MD), and concentrated using an ethanol precipitation protocol with GlycoBlue following manufacturer’s instructions (Thermofisher, Frederick, MD). DNA concentration and integrity were measured by Nanodrop spectrophotometer (Thermofisher). Intact, hypermutated and/or 3’ defective, and 5’ defective HIV-1 copies/million cells were determined by droplet digital PCR (ddPCR) using the IPDA (45), where HIV-1 and human RPP30 reactions were conducted independently in parallel, and copies were normalized to the quantity of input DNA. Briefly, primers and probes have been previously reported (46), and, in each ddPCR reaction, a median 7.5 ng (for RPP30) or a median 750 ng (for HIV-1) of genomic DNA was combined with ddPCR Supermix for Probes (no dUTPs, BioRad), primers (final concentration 900 nM, Integrated DNA Technologies), and probe(s) (final concentration 250 nM, ThermoFisher Scientific). Droplets were analyzed on a QX200 Droplet Reader (BioRad) using QuantaSoft software (BioRad, version 1.7.4), where replicate wells were merged prior to analysis. Four technical replicates were performed for each participant sample. Intact HIV-1 copies (Ψ and *env* double-positive droplets) were corrected for DNA shearing based on the frequency of RPP30 and RPP30-Shear double-positive droplets.

### Mass Cytometry

Mass cytometry was performed in eight individuals treated with ALN and seven who took placebo using an optimized panel of 32 markers.

#### Panel design and conjugation of antibodies

Pairing of metals with antibodies was conducted using Fluidigm software (San Francisco, CA). When commercial antibodies were not available, custom conjugations were performed using MaxPar X8 Antibody Labeling Kit (Fluidigm). Cell populations were phenotypically defined as previously reported (47, 48) and described in Supplementary Table S9.

#### Sample Preparation

In each experiment, 3×10^6^ PBMCs (pre-stained with CD45/89Y) were combined with 5×10^5^ spiked control PBMCs from a single HIV-1-infected control donor that was consistently used in all the experiments (pre-stained with CD45/115In), as described (49). Prior to 30min RT incubation with surface marker antibodies in Cell Staining Buffer (CSB, Fluidigm), cell suspensions were viability-stained using monoisotopic Cell-ID Cisplatin 198 (Fluidigm) following manufacturers’ recommendations. Cells were washed twice in CSB, fixed for 60 min with 2% Paraformaldehyde (Electron Microscopy Sciences) in PBS (Rockland Immunochemicals), and washed once with 1x Perm/Wash Buffer (PWB, BioLegend). Cells were incubated for 30 min at 37°C with the intracellular antibody cocktail in PWB. Stained cell suspensions were barcoded (20-Plex Pd Barcoding Kit, Fluidigm), pooled, resuspended in 62.5 nM Cell-ID Intercalator-Ir (Fluidigm) in Maxpar Fix and Perm Buffer (Fluidigm) and incubated overnight at 4°C. Immediately prior to data acquisition, samples were centrifuged, washed in CSB, followed by a wash in Cell Acquisition Solution (CAS, Fluidigm), filtered (40 micron Cell Strainer, FlowMi), and cell concentration adjusted in CAS to 0.5×10^6^ cells/mL with addition of 10% (v/v) EQ Four Element Calibration Beads (Fluidigm).

#### Single cell data acquisition

Data were acquired on a Helios model mass cytometer (Fluidigm) equipped with a WB injector.

#### Single cell data analysis

Data files were bead normalized and debarcoded using software provided by Fluidigm. Analysis was performed in collaboration with Astrolabe Diagnostics (50). Single-cell data was clustered using the FlowSOM R package (50, 51). Differential abundance analysis was done using the edgeR package (52, 53) following a previously reported method (54). Data is presented as relative frequency of cell populations related to the total number of alive events acquired. Cluster labeling, method implementation, and visualization were done through the Astrolabe Cytometry Platform (Astrolabe Diagnostics, Inc.).

### HIV-1-specific T cell responses

ELISPOT was performed as previously reported (55). Briefly, cryopreserved PBMCs from participants treated *in vivo* with ALN or PLACEBO were thawed and rested overnight then added to ELISpot plates (Merck, Millipore) in quadruplicate (1 or 1.25 × 10^5^ PBMC/ well) with peptides pools of optimal CD8 T cell epitopes from either HIV-1-1 Gag/Nef (n=109, 2µg/ml, CTL-A) or Influenza, HCMV, and Epstein-Barr Viruses (n= 55, 2µg/ml, FEC55). PHA positive controls and media-only controls were also run. Samples were counted independently three times, and cell counts were averaged resulting in <10% variation. No minimum cell viability or cell recovery criteria was applied. Timepoints for each participant were run together. Antigens in thawed peptide plates were mixed with 1:1 with PBMCs in the ELISpot plate to a final concentration of 2 μg/ml and incubated for 18-20 hours at 37°C, 5% CO2. Coating, development (MabTech), and reading of ELISpot plates (AID Reader) has been described previously. Positive T cell responses were defined as ≥30 SFU per million, > 4 times the average of replicate background wells. Zero values were not accepted in any replicate of antigen-stimulated wells.

### Statistics

Nonparametric, two-sided, exact tests were used to make comparisons. A Mann-Whitney U test was used for comparisons between different groups, a Wilcoxon signed-rank test was used for analyzing repeated measures within the same groups, and a Fisher’s exact test was used for comparisons between categorical variables. Values for *ex vivo* analysis of HIV-1 *gag* caRNA below the limit of quantification (LOQ) were assigned a value of half the limit of quantification (LOQ=5 copies/mL). SLDAssay,(56) a software package in R for serial limiting dilution assays, was used to compute maximum-likelihood bias-corrected estimates for IUPM, accompanied by an exact 95% CI and p-value for goodness of fit (PGOF). Tests for trend with increasing time on intervention were performed using a two-sided, Mann-Kendall test for monotonic trend (in either direction) for *in vivo* analyses of both HIV-1 caRNA *pol* copies and HIV-1 *gag* DNA copies/10^5^ PBMC and *in vitro* analyses of IPDA. For the *ex vivo* analysis of TLR2 expression on isolated CD4 T cells, a Friedman test was used to determine if there were differences in modulation of expression at any timepoint 2, 4, 6, and 20 hours post-intervention. For the single cell data analysis from mass cytometry experiments, pairwise comparisons between basal and weeks 2, 24, and 48 post-intervention were performed using a Wilcoxon signed-rank test. Pairwise comparisons for HIV-1 DNA species between baseline and week 2 were performed by a Wilcoxon signed-rank test. P-values were adjusted for multiple comparisons using the Holm-Bonferroni method. P-values between differentially expressed genes in the PAM untreated/treated data set and the CD4 T cell HIV-1 infection data set were calculated using DESeq2, which calculates the Wald test statistic for each gene.

Graphs were created using GraphPad Prism v.9.4 (GraphPad Software, San Diego, CA, USA), and statistical analyses were performed in either GraphPad Prism v.9.4 or R version 3.5.3 (RStudio version 1.2).

### Study Approval

All participants provided written informed consent prior to inclusion in the study, and studies were approved by the George Washington and University of North Carolina Institutional Review Board.

## DISCUSSION

In this work, we show the novel use of N-BPs as LRAs. Based on a careful examination of HIV-1 caRNA and replication-competent virus production after *ex vivo* exposure of isolated rCD4 T cells from PLWH on ART, we demonstrate the capacity of N-BPs to reverse latency. These results were further corroborated using samples from *in vivo* treated PLWH from a previous clinical trial (36). We detected perturbations of the viral reservoir consisting of an induction in HIV-1 caRNA in four participants, and a decay in three others in whom a decay in total HIV-1 DNA was also observed. However, limited sample size and cell availability precluded providing evidence of an impact on the viral reservoir. Despite these limitations, we show that N-BPs exert a potent induction of viral replication, a finding that warrants further clinical investigation.

HIV-1 caRNA reflects the size of the HIV-1 reservoir (38, 57) since it represents intracellular transcripts that originate from latently infected cells in virally suppressed PLWH (37, 38). Confirming previous studies (58, 59), women on ART had lower frequency of infection measured by HIV-1 caRNA. ALN induced perturbations on HIV-1 caRNA in 78% of the participants which included both increased and decreased levels upon intervention, highlighting the complexity of caRNA dynamics. Variability of the HIV-1 caRNA slopes in our study may reflect when and for how long the LRA effect is measurable after once a week ALN administration. Encouragingly, in three of the ALN-treated participants changes in HIV-1 caRNA coincided with a decay in total HIV-1 DNA levels. Similarly, a recent study also observed variable HIV-1 caRNA slopes with increases and decreases after treatment with a broadly neutralizing antibody and VOR, although no changes in DNA were detected (39).

Previous studies have shown decay for intact but not defective HIV-1 DNA species in PLWH on ART (60–62). Due to sample limitations, our study used total PBMCs for IPDA. We found that levels of intact provirus measured in PBMC were similar to previous reports and correlated with total HIV-DNA, as previously reported (46, 59–61, 63–65). Participants in our small study cohort had received ART for a mean of 4.4 years and changes in intact, total or defective HIV-1 species did not associate with time of exposure to ART. Overall, levels of HIV-1 DNA species were comparable between ALN-treated and placebo groups throughout the study. We did however detect a trend towards a decay from baseline to week two in the ALN group in intact provirus. By contrast, we found no evidence of decay either in total and defective HIV-1 DNA in the placebo group. Overall, while our results are encouraging, they need to be interpreted cautiously due to our small sample size, the inherent variability of the assay and complex reservoir dynamics, especially during the initial years of ART exposure. Therefore, the potential effect of ALN on the viral reservoir warrants further investigation using a larger cohort with the specific objective of analyzing the novel use of ALN as an LRA and immunomodulatory agent.

Treatment with N-BPs induces activation of Vδ2 T cells (12–14) and a prior study showed induction of HIV-specific T cell responses after *in vivo* treatment with Zol and IL-2 (66) and our previous studies demonstrated the *ex vivo* capacity of Vδ2 T cells to clear reactivated latently HIV-infected cells (18, 19). *In vivo* administration of ALN activated both Vδ2 T cells and CD8 T cells to produce GzmB, showing that N-BP administration additionally enhance cellular immunity although the effect on the viral reservoir was not evident. Previous studies analyzing the effect of N-BPs on overall circulating immune cell populations are scarce. Intravenous PAM treatment *in vivo* induced transient decreases in the frequency of circulating T cells (67). The longitudinal decay of EM-CD8 T cells in our study may be consistent with redistribution to tissue sites, although preclinical models will be required to confirm this possibility. A detailed immunological profiling of the eight participants suggests that prior host immunity, i.e., constant Vδ2 T cell frequency, sustained EM-CD8 T cells and increased GzmB production over time, may play a role eliminating the persistent viral reservoir.

We propose a model whereby N-BPs induce both HIV-1 reactivation and direct activation of Vδ2 T cells’ effector functions, that in turn enhance cytotoxic properties from other effector cells (20, 21, 68, 69). For the reactivation effect, although further investigation is required to understand how N-BPs reactivate latent HIV, our results suggest that c-Jun, but not NFκB signaling, leads to viral reactivation. Increased phosphorylation at the Ser73 residue of c-Jun (35) after exposure to Zol, provides evidence of the involvement of this pathway on viral reactivation (33, 34). In our experiments, cells were exposed to one single N-BPs dose for 15 minutes and only one phosphorylation site was analyzed. In addition, since Zol has been reported to be the most potent N-BP (70), *in vitro* exposure to PAM or ALN may require higher doses or longer times to detect an effect in c-Jun phosphorylation, and possibly other residues and/or AP-1 components.

One of the advantages of investigating FDA-approved drugs for use in other indications includes the known safety and toxicity profiles. ALN’s safety profile is well known and, although there are side effects that need to be carefully monitored, these drugs are currently used as first-line treatment for osteoporosis (31). We have discovered that N-BPs are a novel class of potent LRAs that induce both reactivation of persistent HIV-1 and effector cellular functions in PLWH on suppressive ART. We present the first evidence in humans that treatment with N-BPs may be further pursued in the context of HIV cure, possibly in combination with other interventions aimed at enhancing pre-existing immunity.

## ACKNOWLEDGEMENTS

Authors would like to thank all participants who voluntarily donate their samples and contribute to finding a cure for HIV. Dr. Siegel and Ms. Langlands for clinical support at GWU, Dr. Margolis and Ms. Kuruc at UNC. Dr. Archin, Ms. Allard and Ms. James for technical support. Dr. Amir (Astrolabe Diagnostics) for mass cytometry data analysis and critical reading the paper. Dr. Chiapinelli for feedback on RNA-seq data. Drs. Eron, Margolis, Archin and Overton (on behalf of the ACTG) read the manuscript. We thank the ACTG for kindly providing the ACTG A5163 clinical samples, and Dr. Eron who facilitated obtaining the samples.

## AUTHOR CONTRIBUTIONS

Conception of the study: NSS. Data collection: MS, MLC, BTM, MAI, CPW, AC, CG, JK, SS, YX, DC. Data analysis and interpretation: MS, AMKW, ARW, MLC, SGR, SRS, MAI, CPW, ARC, YHT, YX, MGH, AB, BRJ, JSP, NG, NSS. Drafting the article: MS, AMKW, SGR, SRS, BTM, AW, MGH, NG, NSS. Critical revision of the article: MS, AMKW, ARW, JK, ARC, CG, AB, MGH, JSP, NG. Final approval of the version to be published: MS, AMKW, ARW, SGR, BTM, AW, AB, BRJ, JSP, MGH, NG, NSS.

## ADDITIONAL INFORMATION

Supplementary Information is available for this paper

Further information and requests for resources and reagents should be directed to and will be fulfilled by Natalia Soriano-Sarabia.

## Supplementary Material

**Supplementary Table 1.**
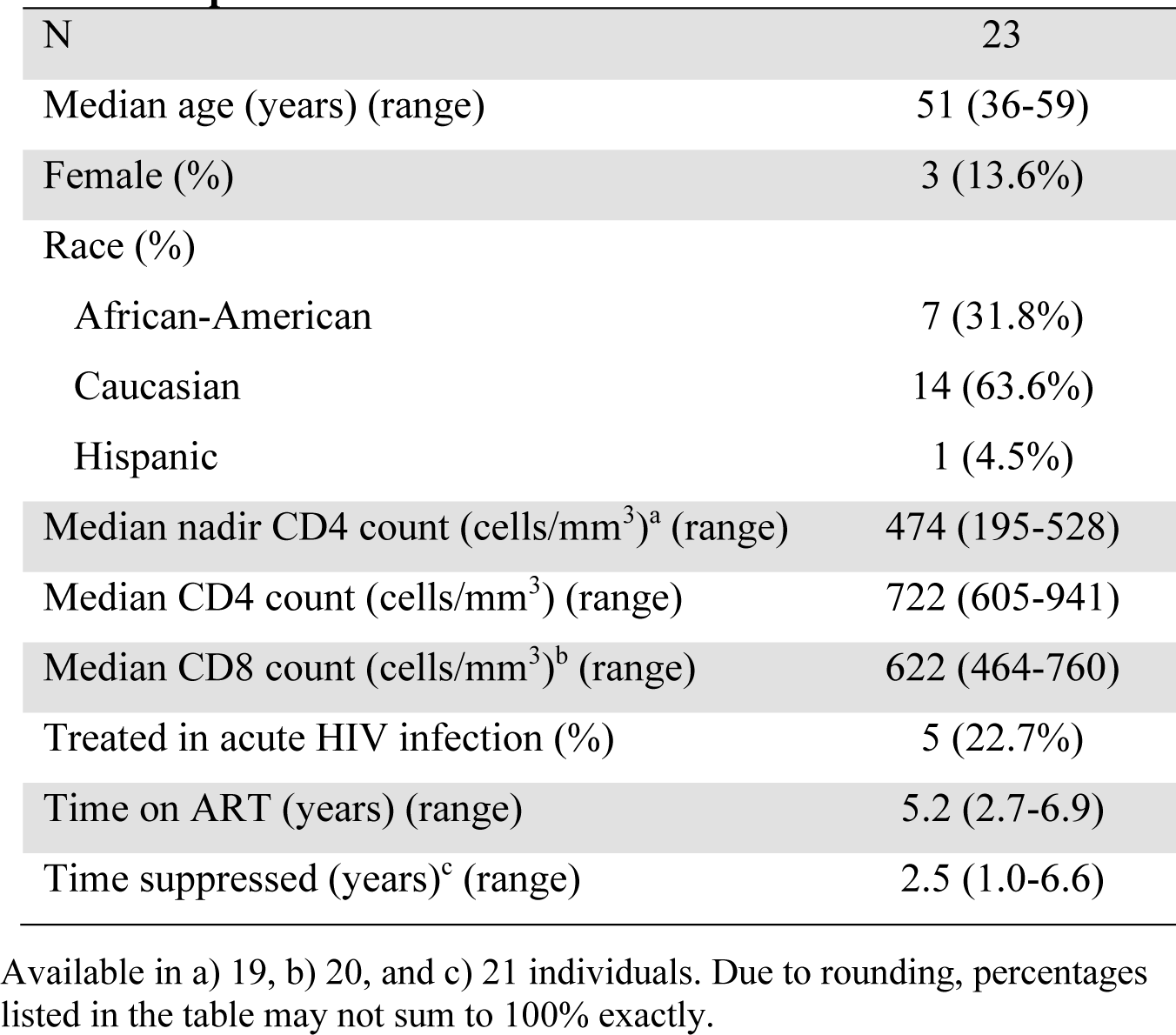
Characteristics of PLWH included in *ex vivo* experiments.

|  |  |
| --- | --- |
| N | 23 |
| Median age (years) (range) | 51 (36-59) |
| Female (%) | 3 (13.6%) |
| Race (%) |  |
| African-American | 7 (31.8%) |
| Caucasian | 14 (63.6%) |
| Hispanic | 1 (4.5%) |
| Median nadir CD4 count (cells/mm<sup>3</sup>)<sup>a</sup> (range) | 474 (195-528) |
| Median CD4 count (cells/mm<sup>3</sup>) (range) | 722 (605-941) |
| Median CD8 count (cells/mm<sup>3</sup>)<sup>b</sup> (range) | 622 (464-760) |
| Treated in acute HIV infection (%) | 5 (22.7%) |
| Time on ART (years) (range) | 5.2 (2.7-6.9) |
| Time suppressed (years)<sup>c</sup> (range) | 2.5 (1.0-6.6) |

Available in a) 19, b) 20, and c) 21 individuals. Due to rounding, percentages listed in the table may not sum to 100% exactly.

**Supplementary Table 4.** Individual clinical trial participant characteristics.

| Group | Pt. | Sex | Race/Ethnicity | Age | CD4 counts | CD8 counts | CD4/CD8 ratio | Years on ART |
| --- | --- | --- | --- | --- | --- | --- | --- | --- |
| ALN | 101 | F | Black Non-Hispanic | 48 | 974 | 669 | 1.46 | 1.46 |
| ALN | 102 | F | Black Non-Hispanic | 43 | 492 | 1132 | 0.43 | 4.74 |
| ALN | 103 | M | White Non-Hispanic | 44 | 679 | 868 | 0.78 | 9.53 |
| ALN | 104 | F | White Non-Hispanic | 39 | 687 | 1061 | 0.65 | 5.49 |
| ALN | 105 | M | White Non-Hispanic | 55 | 408 | 957 | 0.43 | 2.18 |
| ALN | 106 | F | More than one race | 33 | 153 | 484 | 0.32 | 0.63 |
| ALN | 107 | M | White Non-Hispanic | 48 | 429 | 891 | 0.48 | 5.3 |
| ALN | 108 | M | White Non-Hispanic | 58 | 1290 | 1935 | 0.67 | 1.44 |
| ALN | 109 | F | Hispanic (Regardless of Race) | 48 | 257 | 971 | 0.26 | 2.24 |
| ALN | 110 | F | Black Non-Hispanic | 49 | 232 | 506 | 0.46 | 4.93 |
| ALN | 111 | M | White Non-Hispanic | 62 | 1216 | 1687 | 0.72 | 1.9 |
| ALN | 112 | F | Asian, Pacific Islander | 54 | 219 | 804 | 0.27 | 0.51 |
| ALN | 113 | M | White Non-Hispanic | 47 | 559 | 505 | 1.11 | 0.19 |
| ALN | 114 | M | Black Non-Hispanic | 56 | 307 | 725 | 0.42 | 1.26 |
| ALN | 115 | M | White Non-Hispanic | 54 | 876 | 1169 | 0.75 | 16.8 |
| ALN | 116 | M | Black Non-Hispanic | 48 | 864 | 1620 | 0.53 | 2.36 |
| ALN | 117 | M | White Non-Hispanic | 49 | 464 | 1824 | 0.25 | 5.06 |
| ALN | 118 | M | Asian, Pacific Islander | 52 | 105 | 1328 | 0.08 | 1.6 |
| ALN | 119 | F | Asian, Pacific Islander | 52 | 664 | 708 | 0.94 | 4.31 |
| ALN | 120 | M | White Non-Hispanic | 47 | 269 | 1366 | 0.20 | 0.83 |
| ALN | 121 | F | Hispanic (Regardless of Race) | 60 | 906 | 1324 | 0.68 | 1.84 |
| ALN | 122 | M | White Non-Hispanic | 42 | 1117 | 549 | 2.03 | 0.93 |
| ALN | 123 | M | White Non-Hispanic | 63 | 262 | 759 | 0.35 | 0.85 |
| **Median** |  |  |  | 49 | 492 | 957 | 0.48 | 1.9 |
| Placebo | 201 | M | Black Non-Hispanic | 40 | 653 | 1448 | 0.45 | 2.06 |
| Placebo | 202 | M | Hispanic (Regardless of Race) | 55 | 491 | 1601 | 0.31 | 0.57 |
| Placebo | 203 | M | White Non-Hispanic | 53 | 412 | 1510 | 0.27 | 2.01 |
| Placebo | 204 | F | White Non-Hispanic | 33 | 460 | 841 | 0.55 | 1.75 |
| Placebo | 205 | F | Black Non-Hispanic | 36 | 638 | 723 | 0.88 | 1.29 |
| Placebo | 206 | M | White Non-Hispanic | 42 | 483 | 372 | 1.30 | 5.07 |
| Placebo | 207 | M | White Non-Hispanic | 30 | 356 | 1449 | 0.25 | 0.72 |
| Placebo | 208 | M | White Non-Hispanic | 56 | 416 | 814 | 0.51 | 3.11 |
| Placebo | 209 | M | White Non-Hispanic | 57 | 1003 | 1405 | 0.71 | 4.72 |
| Placebo | 210 | F | White Non-Hispanic | 58 | 556 | 391 | 1.42 | 4.4 |
| Placebo | 211 | F | Black Non-Hispanic | 60 | 396 | 1328 | 0.30 | 0.83 |
| Placebo | 212 | M | Hispanic (Regardless of Race) | 45 | 221 | 488 | 0.45 | 7.31 |
| Placebo | 213 | M | White Non-Hispanic | 35 | 212 | 570 | 0.37 | 0.86 |
| Placebo | 214 | M | White Non-Hispanic | 43 | 493 | 775 | 0.64 | 9.88 |
| Placebo | 215 | F | Hispanic (Regardless of Race) | 45 | 465 | 581 | 0.80 | 3 |
| Placebo | 216 | M | White Non-Hispanic | 55 | 375 | 666 | 0.56 | 1.04 |
| Placebo | 217 | F | White Non-Hispanic | 68 | 189 | 192 | 0.98 | 5.13 |
| Placebo | 218 | M | White Non-Hispanic | 35 | 739 | 1443 | 0.51 | 2.13 |
| Placebo | 219 | F | Black Non-Hispanic | 41 | 810 | 718 | 1.13 | 8.16 |
| Placebo | 220 | M | White Non-Hispanic | 44 | 327 | 785 | 0.42 | 7.33 |
| Placebo | 221 | M | White Non-Hispanic | 41 | 1037 | 754 | 1.38 | 5.31 |
| **Median** |  |  |  | 44 | 465 | 775 | 0.55 | 3 |

Grey shaded samples were used for IPDA.

**Supplementary Table 5.** IPDA participant characteristics

|  | **ALN (n=9)** | **Placebo (n=7)** | **p-value<sup>a</sup>** |
| --- | --- | --- | --- |
| Biological sex, No. (%) |  |  | 0.173 |
| Female | 1 (11.1%) | 3 (42.9%) |  |
| Male | 8 (88.9%) | 4 (57.1%) |  |
| Race/Ethnicity, No. (%) |  |  | **0.033** |
| White non-Hispanic | 5 (55.6%) | 4 (57.1%) |  |
| Black non-Hispanic | 2 (22.2%) | 1 (14.3%) |  |
| Hispanic (regardless of race) | 0 (0.0%) | 2 (28.6%) |  |
| Asian, Pacific Islander | 2 (22.2%) | 0 (0.0%) |  |
| Age (years)<sup>b</sup> | 49 (47 - 56) | 45 (35 - 68) | 0.596 |
| CD4 count (cells/mm<sup>3</sup>)<sup>b</sup> | 464 (105 - 876) | 375 (189 - 493) | 0.204 |
| CD8 count (cells/mm<sup>3</sup>)<sup>b</sup> | 1169 (505 - 1824) | 581 (192 - 1328) | **0.039** |
| CD4/CD8 ratio<sup>b</sup> | 0.48 (0.08 - 1.11) | 0.56 (0.30 - 0.98) | 0.597 |
| Years on ART<sup>b</sup> | 2.36 (0.19 - 16.75) | 3.00 (0.83 - 9.88) | 0.691 |

<sup>a</sup>P-values are calculated using Fisher's exact test (categorical variables) or Mann-Whitney U test (continuous variables). <sup>b</sup>Median and (range) are presented.

**Supplementary Table 6.** Characteristics of participants included in caRNA and mass cytometry assays

|  | **ALN (n=15)** | **Placebo (n=14)** | **p-value<sup>a</sup>** |
| --- | --- | --- | --- |
| Biological sex, No. (%) |  |  |  |
| Female | 8 (0.53%) | 4 (0.29%) |  |
| Male | 7 (0.47%) | 10 (0.71%) |  |
| Race, No. (%) |  |  |  |
| White non-Hispanic | 8 (0.53%) | 10 (0.71%) |  |
| Black non-Hispanic | 1 (0.07%) | 3 (0.21%) |  |
| Hispanic (regardless of race) | 2 (0.13%) | 1 (0.07%) |  |
| Asian, Pacific Islander | 1 (0.07%) | 0 |  |
| Age (years)<sup>b</sup> | 48 (43-55) | 48 (35-56) | 0.62 |
| CD4 count (cells/mm<sup>3</sup>)<sup>b</sup> | 461 (238-902) | 483 (412-638) | 0.88 |
| CD8 count (cells/mm<sup>3</sup>)<sup>b</sup> | 924 (703-1114) | 1309 (723-1449) | 0.79 |
| Years on ART<sup>b</sup> | 2.5 (1.4-5.2) | 2.0 (1.3-4.7) | 0.69 |

<sup>a</sup>P-values are calculated using Fisher's exact test (categorical variables) or Mann-Whitney U test (continuous variables). <sup>b</sup>Median and (range) are displayed.

**Supplementary Table 7.** Time trend analysis of IPDA measures

| **Supplementary Table 7.** Time trend analysis of IPDA measures |  |  |
| --- | --- | --- |
| **Measure** | **Kendall's Tau** | **p-Value** |
| *ALN - week 0 to 48* |  |  |
| Total HIV DNA (copies/10<sup>6</sup> PBMC) | -0.23 | 0.092 |
| Total defective HIV DNA (copies/10<sup>6</sup> PBMC) | -0.24 | 0.074 |
| 3' defective HIV DNA (copies/10<sup>6</sup> PBMC) | **-0.30** | **0.026** |
| 5' defective HIV DNA (copies/10<sup>6</sup> PBMC) | -0.20 | 0.144 |
| Intact HIV DNA (copies/10<sup>6</sup> PBMC) | -0.05 | 0.735 |
| *Placebo - week 0 to 48* |  |  |
| Total HIV DNA (copies/10<sup>6</sup> PBMC) | -0.24 | 0.127 |
| Total defective HIV DNA (copies/10<sup>6</sup> PBMC) | -0.22 | 0.152 |
| 3' defective HIV DNA (copies/10<sup>6</sup> PBMC) | -0.25 | 0.115 |
| 5' defective HIV DNA (copies/10<sup>6</sup> PBMC) | -0.20 | 0.199 |
| Intact HIV DNA (copies/10<sup>6</sup> PBMC) | -0.11 | 0.482 |

<sup>a</sup>Kendall's tau is a rank-based measure of correlation. <sup>b</sup>p-values calculated by the Mann-Kendall test for trend. Abbreviations: IPDA, intact proviral DNA assay; PBMC, peripheral blood mononuclear cells.

**Supplementary Table 8.** Spearman correlations between changes in IPDA measures from baseline to week 2 with clinical parameters at baseline (ALN group)

| Clinical parameter | $r_s$ | p-value | n |
| --- | --- | --- | --- |
| *Intact HIV DNA change<sup>a</sup> correlation* |  |  |  |
| CD4 count (cells/mm<sup>3</sup>) | 0.57 | 0.180 | 7 |
| CD8 count (cells/mm<sup>3</sup>) | -0.71 | 0.071 | 7 |
| CD4/CD8 ratio | 0.71 | 0.071 | 7 |
| Years on ART | -0.04 | 0.939 | 7 |
| Age | 0.35 | 0.443 | 7 |
| *Defective HIV DNA change correlation* |  |  |  |
| CD4 count (cells/mm<sup>3</sup>) | 0.50 | 0.171 | 9 |
| CD8 count (cells/mm<sup>3</sup>) | -0.35 | 0.356 | 9 |
| CD4/CD8 ratio | 0.62 | 0.077 | 9 |
| Years on ART | 0.30 | 0.433 | 9 |
| Age | -0.09 | 0.812 | 9 |
| *Total HIV DNA change correlation* |  |  |  |
| CD4 count (cells/mm<sup>3</sup>) | 0.50 | 0.171 | 9 |
| CD8 count (cells/mm<sup>3</sup>) | -0.35 | 0.356 | 9 |
| CD4/CD8 ratio | 0.62 | 0.077 | 9 |
| Years on ART | 0.30 | 0.433 | 9 |
| Age | -0.09 | 0.812 | 9 |

<sup>a</sup>Change from 0 intact copies to 0 intact copies excluded from analysis
Abbreviations: IPDA, intact proviral DNA assay; ALN, alendronate; ART, antiretroviral therapy.

**Supplementary Table S9.** Phenotypic characterization of immune cell populations for Mass Cytometry.

| Cell subset | Description |
| --- | --- |
| Vδ2 T cell | CD3+ CD19- CD33-, TCRvd1- TCRvd2+ |
| Vδ1 T cell | CD3+ CD19- CD33-, TCRvd1+ TCRvd2- |
| CD8+ T cells (naïve) | CD3+ CD19- CD33-, TCRvd1- TCRvd2-, CD4- CD8+ CD56-, CD45RA+, CCR7+ |
| CD8+ T cells (Effector Memory) | CD3+ CD19- CD33-, TCRvd1- TCRvd2-, CD4- CD8+ CD56-, CD45RA-, CCR7- |
| CD8+ T cells (Central Memory) | CD3+ CD19- CD33-, TCRvd1- TCRvd2-, CD4- CD8+ CD56-, CD45RA-, CCR7+ |
| CD4+ T cells (naïve) | CD3+ CD19- CD33-, TCRvd1- TCRvd2-, CD4+ CD8- CD56-, CD45RA+, CCR7+ |
| CD4+ T cells (Effector Memory) | CD3+ CD19- CD33-, TCRvd1- TCRvd2-, CD4+ CD8- CD56-, CD45RA-, CCR7- |
| CD4+ T cells (Central Memory) | CD3+ CD19- CD33-, TCRvd1- TCRvd2-, CD4+ CD8- CD56-, CD45RA-, CCR7+ |
| NK cells (CD56+ CD16-) | CD3- CD19- CD33-, CD56+ CD123- HLADR-, CD16- |
| NK cells (CD56+ CD16+) | CD3- CD19- CD33-, CD56+ CD123- HLADR-, CD16+ |
| Dendritic cells | CD3- CD19- CD33+, CD14-, CD16- |
| Plasmacytoid dendritic cells | CD3- CD19- CD33-, CD11c- CD123+ HLADR+ |
| Monocytes | CD3- CD19- CD33+, CD14+, CD16- |
| B cells | CD3- CD19+ CD33- |

**Figure S1:**
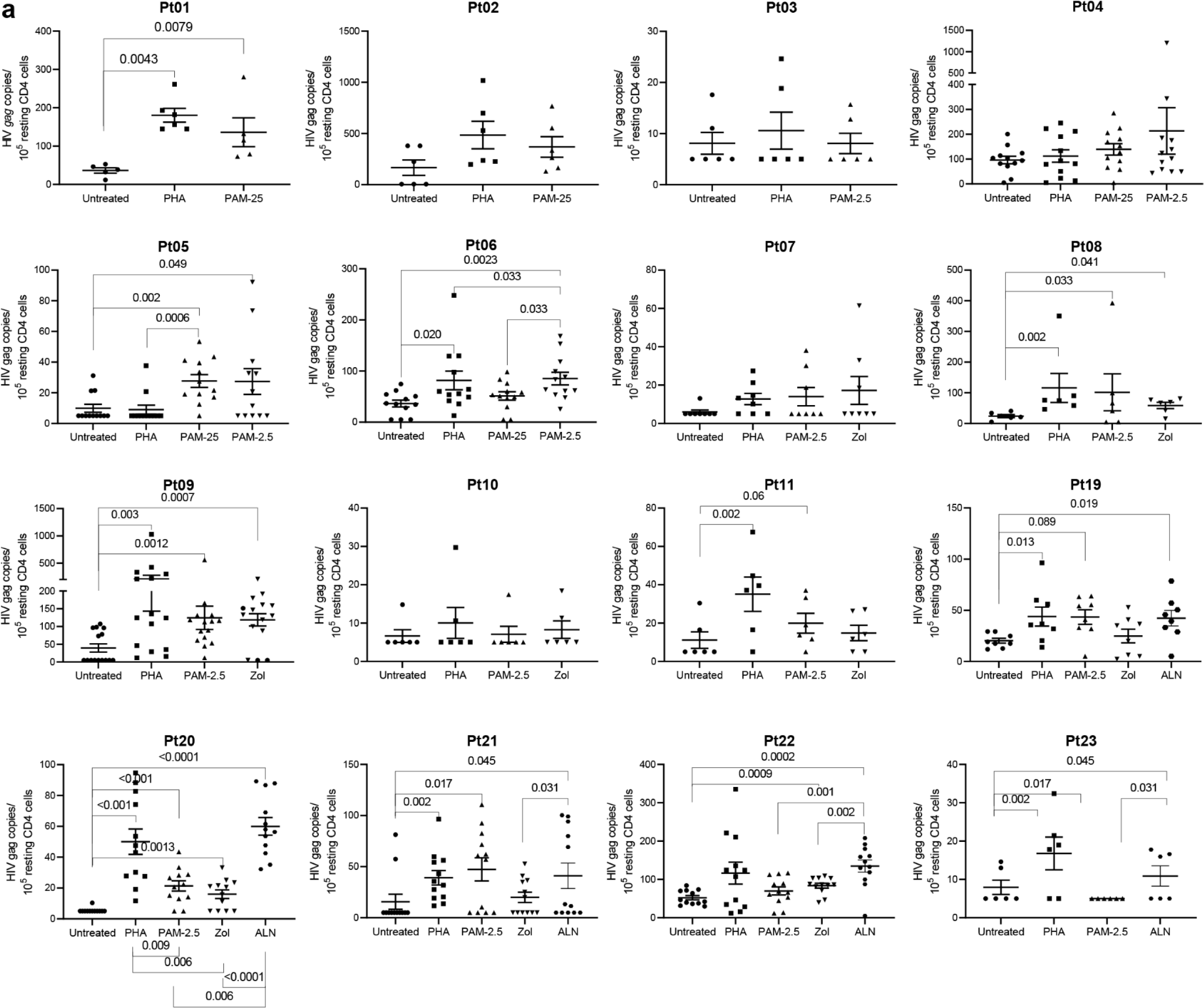
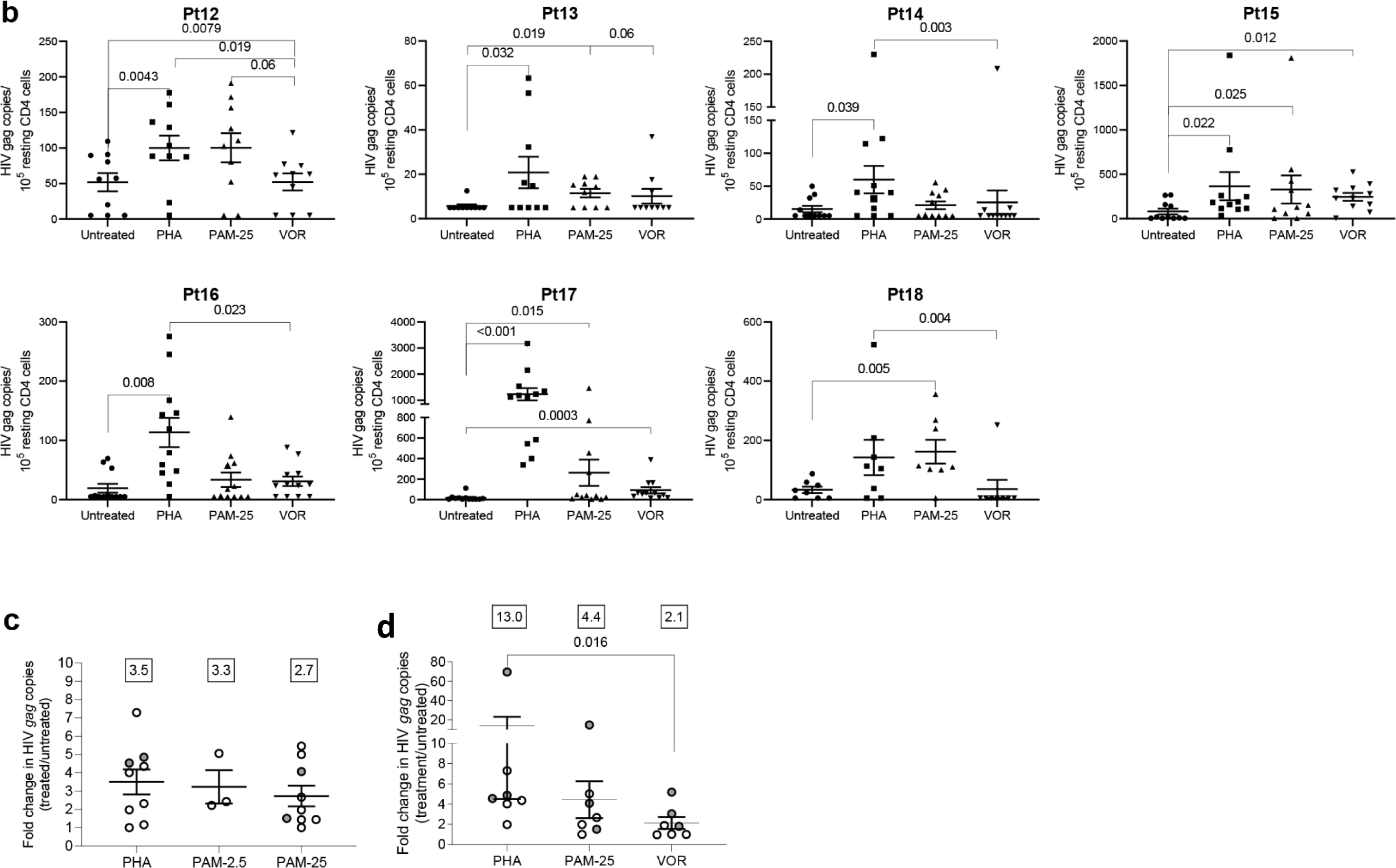
HIV-1 caRNA levels in 23 PLWH on suppressive ART. Individual plots show HIV-1 *gag* caRNA copies/mL in the different conditions tested in each experiment depending on cell availability. **A)** Comparison of PAM, Zol and ALN with PHA. **B)** In Pt12 through Pt18, the capacity of N-BPs to reactivate latent HIV was compared to 500nM VOR. Each graph represents one participant, and each symbol represents one biological replicate of 1×10^6^ isolated rCD4 T cells. Comparison of HIV gag copies induction between **C)** PAM at 2.5µg/mL and 25µg/mL) and **D)** PAM at 25µg/mL and VOR. Pt, participant. P-values were calculated by a Wilcoxon signed-rank test, and only p-values < 0.05 are displayed.

**Figure S2:**
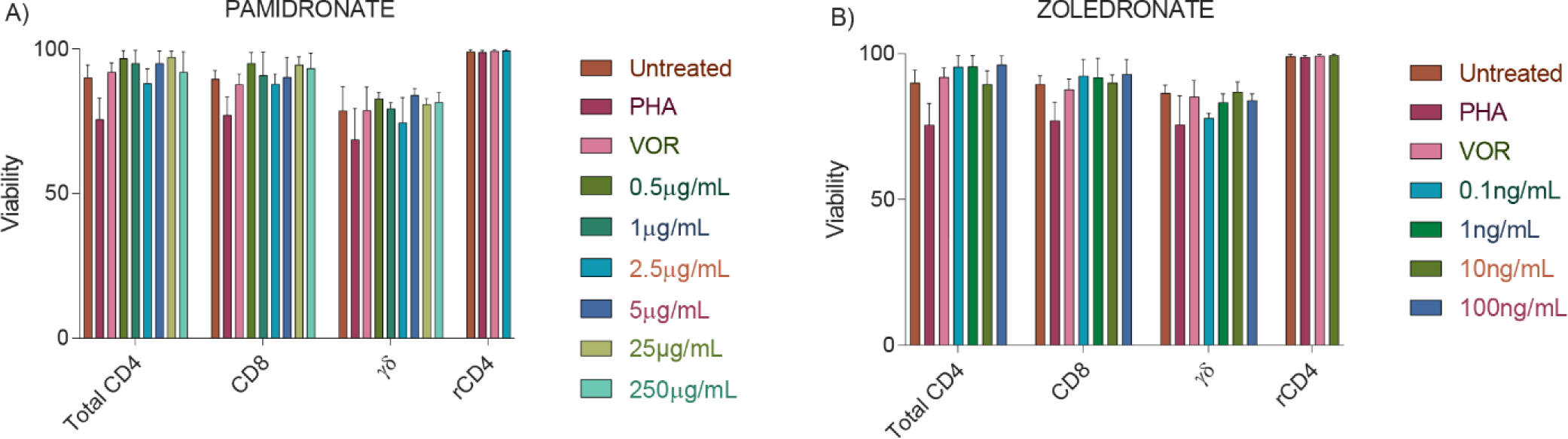
*Ex vivo* viability upon treatment with PAM and Zol. PBMCs from PLWH on suppressed ART were exposed to 2 μg/mL PHA+100 U/mL IL-2 or different concentrations of **A)** PAM or **B)** Zol as specified, and viability (7-AAD) measured by flow cytometry in total CD4 T cells, CD8 T cells, γδ T cells, and resting CD4 (rCD4) T cells. The means of repeated experiments in three to six individuals are presented.

**Figure S3:**
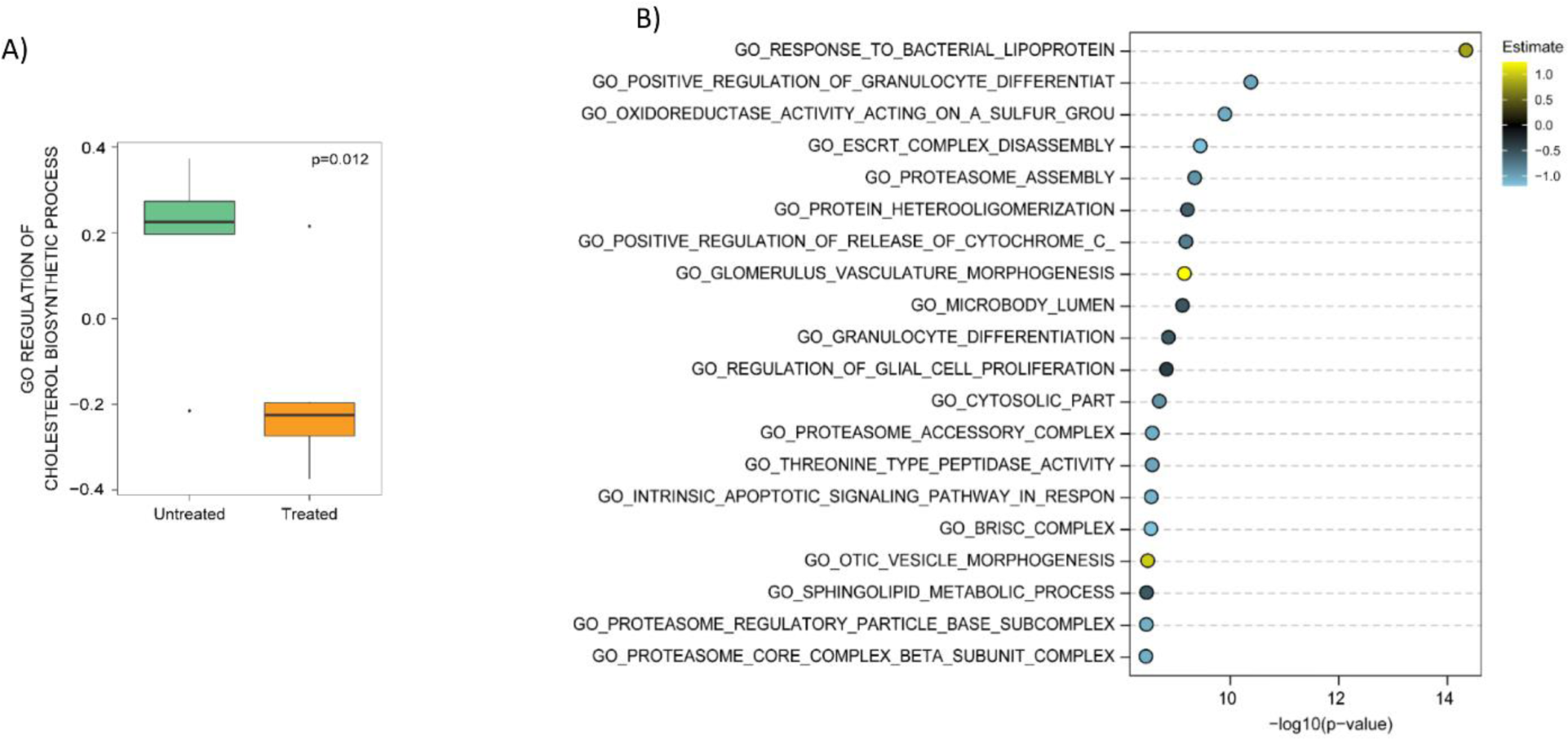
**A)** Boxplot of enrichment score derived from gene set variation analysis (GSVA) of GO annotation of Cholesterol Biosynthesis Regulation comparing untreated and PAM-treated individuals (p=0.012). **B)** Top 20 GO annotations score derived from gene set variation analysis (GSVA) of GO annotation between the untreated and PAM-treated samples. Estimates (coefficients of the predictor treatment) are sorted (on the x-axis) by -log10(p-value) in decreasing direction and color-coded based on the value of the estimate. Yellow: positive estimates; Blue: negative estimates.

**Figure S4:**
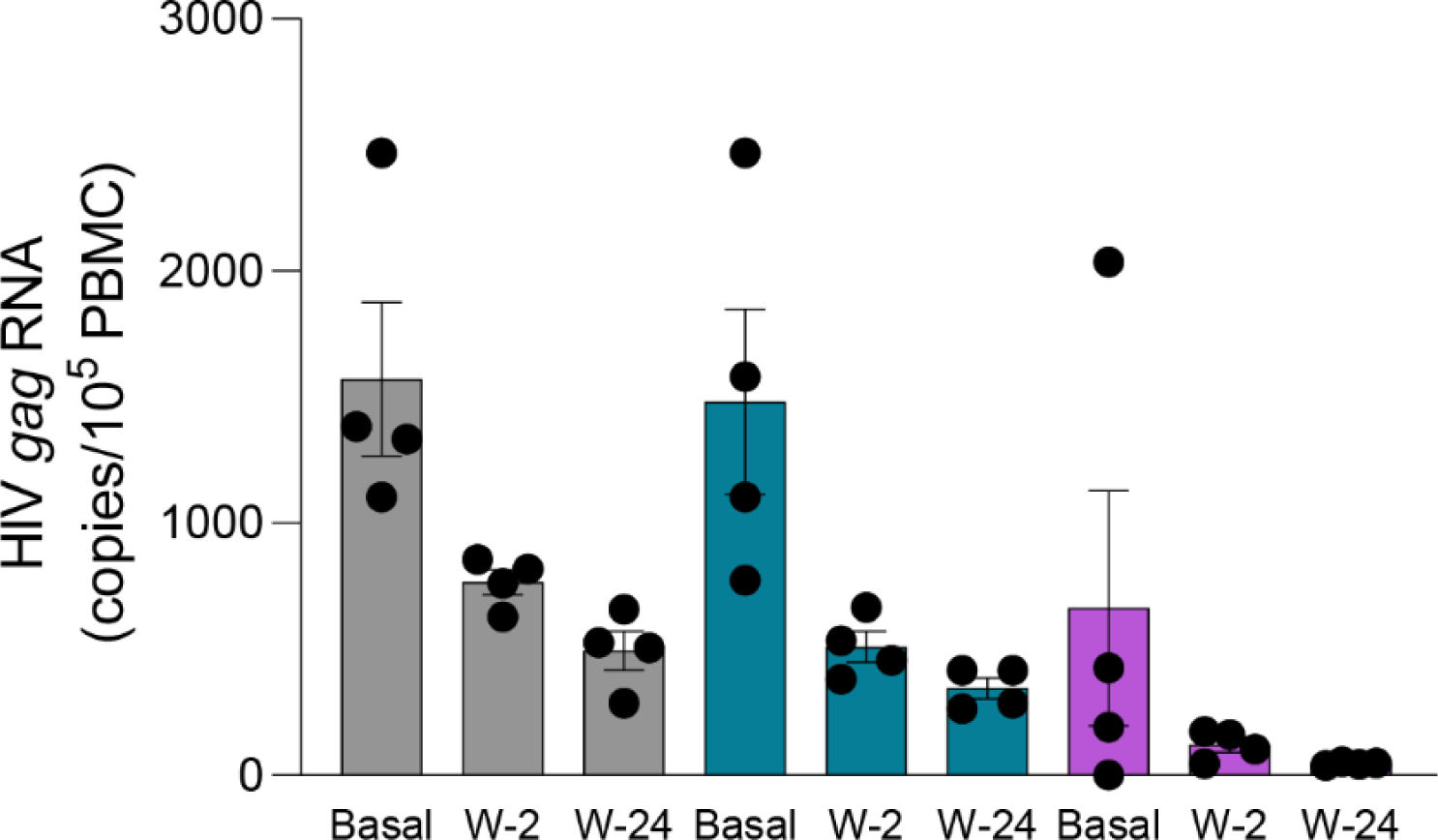
HIV DNA levels in participant 107 at baseline and after ALN treatment. Mean± SEM is represented. Hypermutated 3’defective (grey), 5’ defective (blue) and intact provirus (purple) are presented.

**Figure S5:**
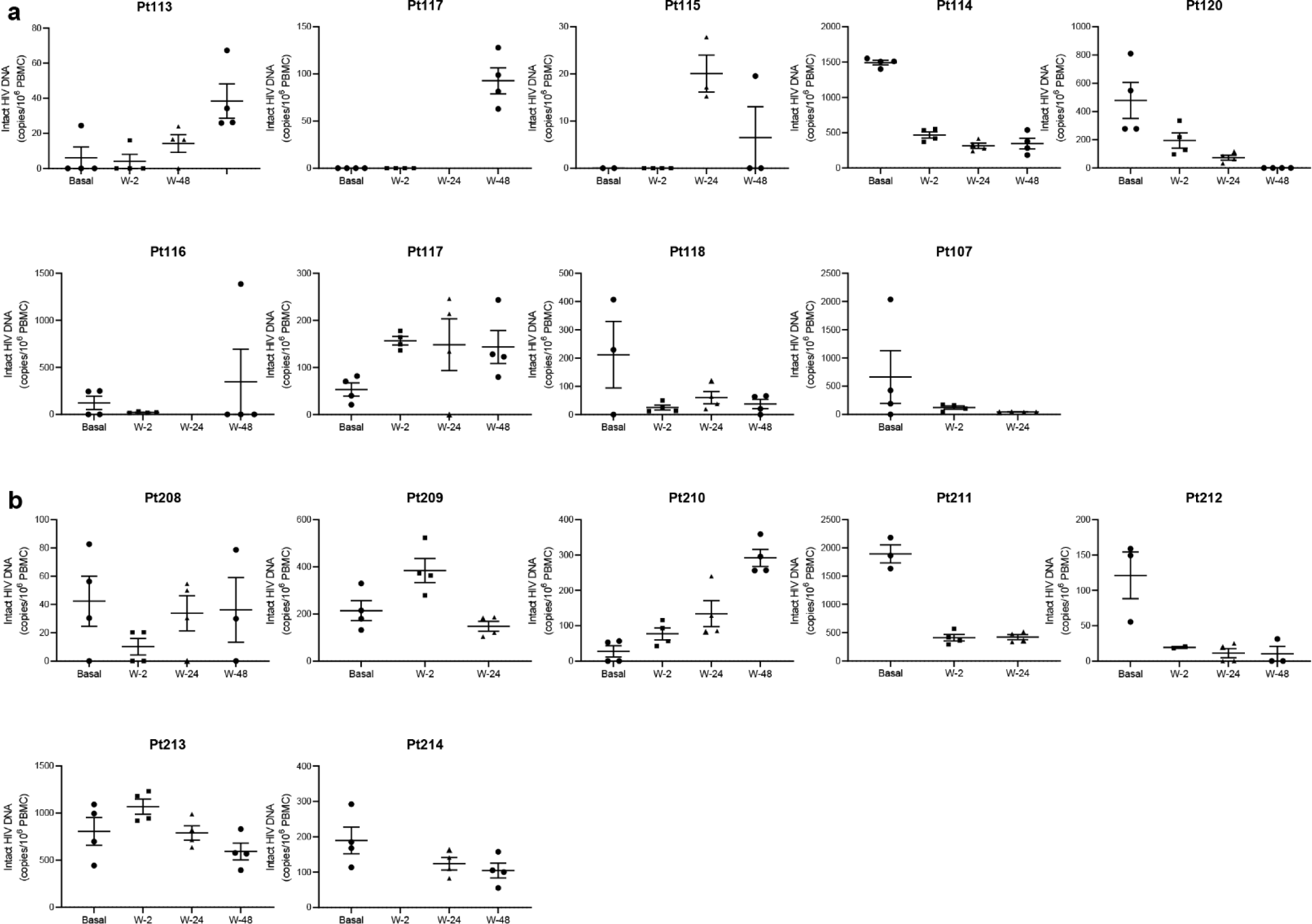
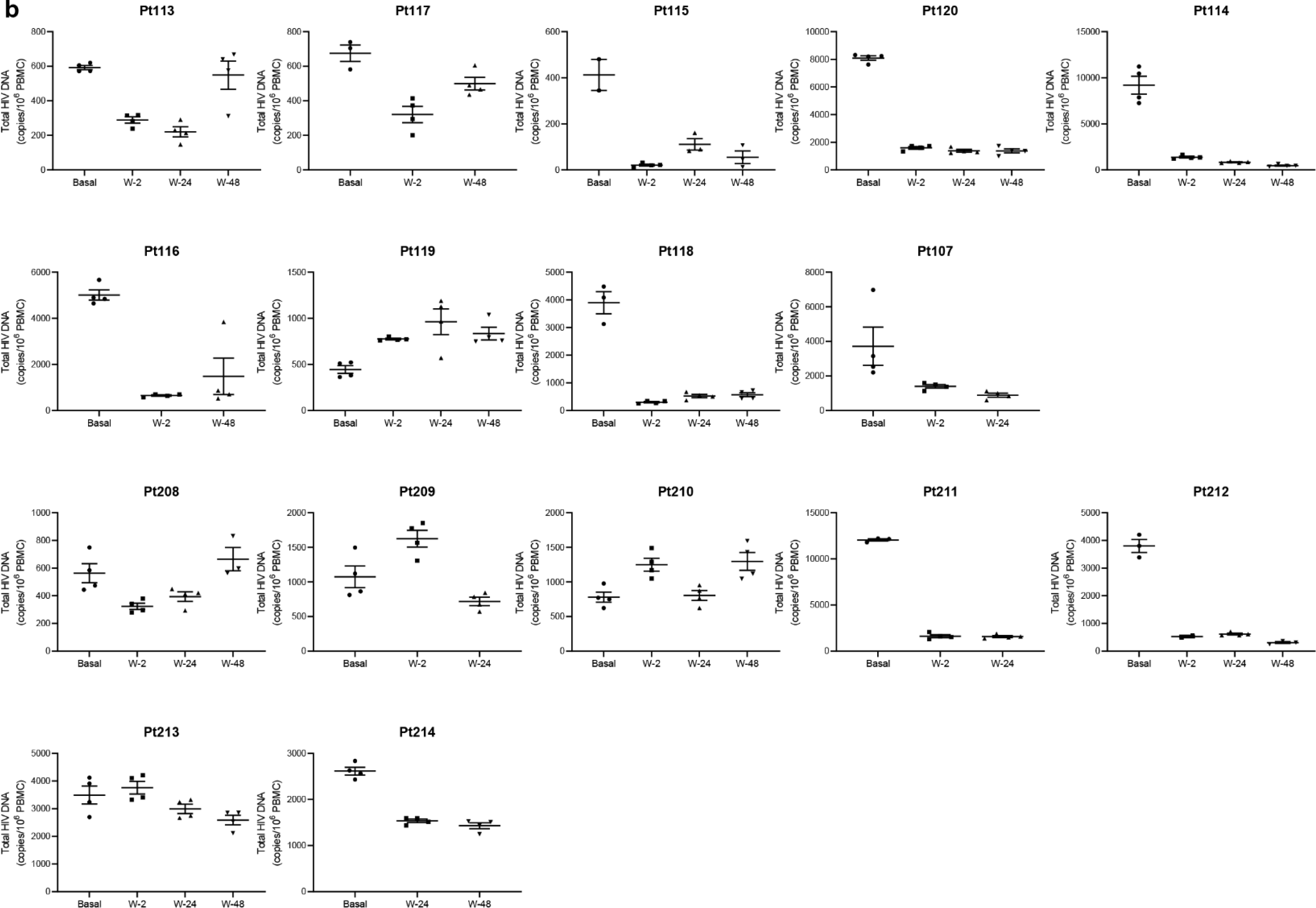
Individual IPDA data showing intact provirus in **A)** ALN and **B)** placebo groups, and total HIV-DNA in **C)** ALN and **D)** placebo groups.

**Figure S6:**
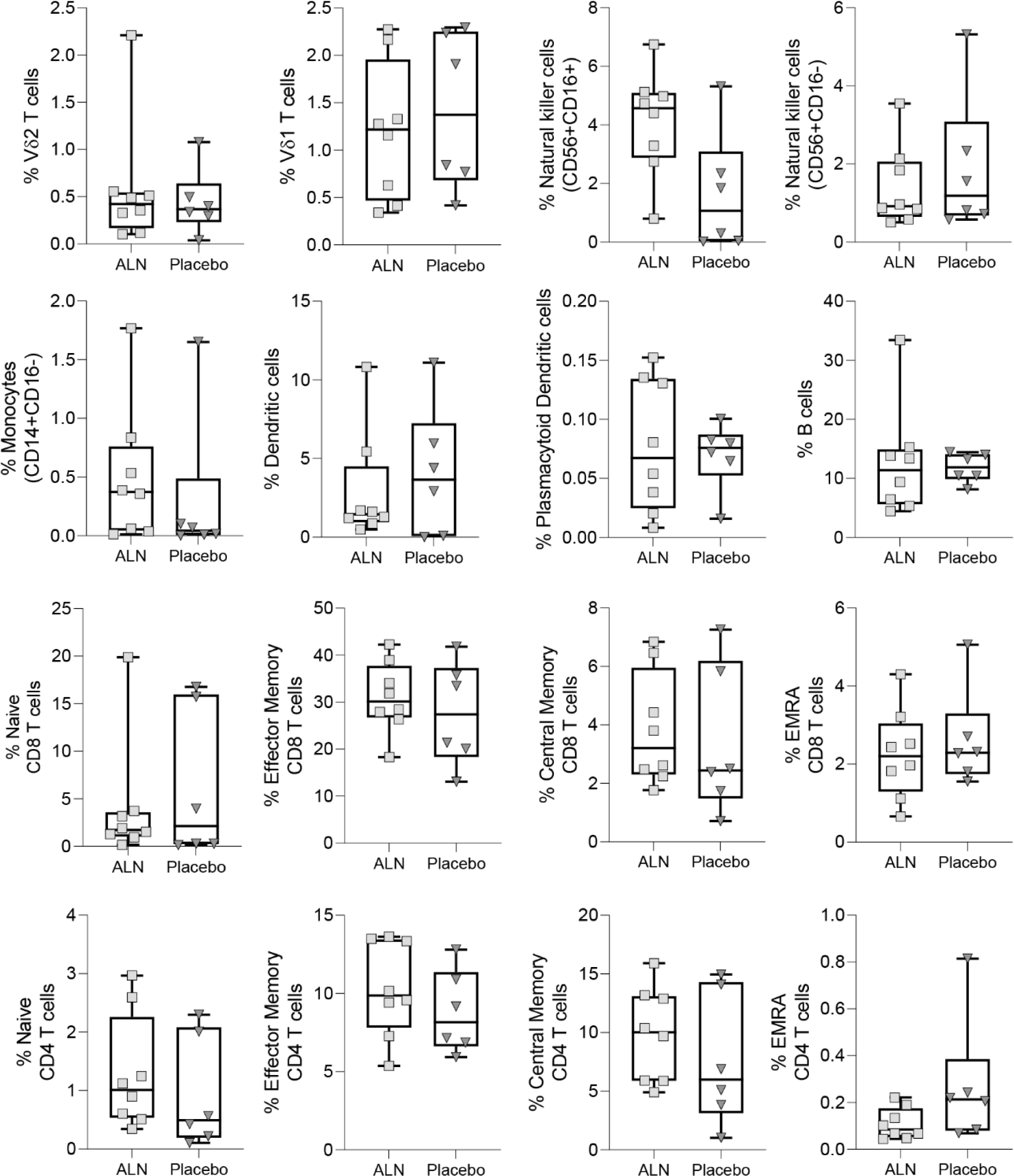
Comparison of basal cell frequencies between individuals who took ALN and placebo. Mass cytometry performed on eight individuals from the ALN group and six from the placebo group. Boxplots display first quartile, median, and third quartile with whiskers range from the minimum to maximum values. Mann-Whitney U test p-values>0.05 for all comparisons between ALN and placebo

**Figure S7:**
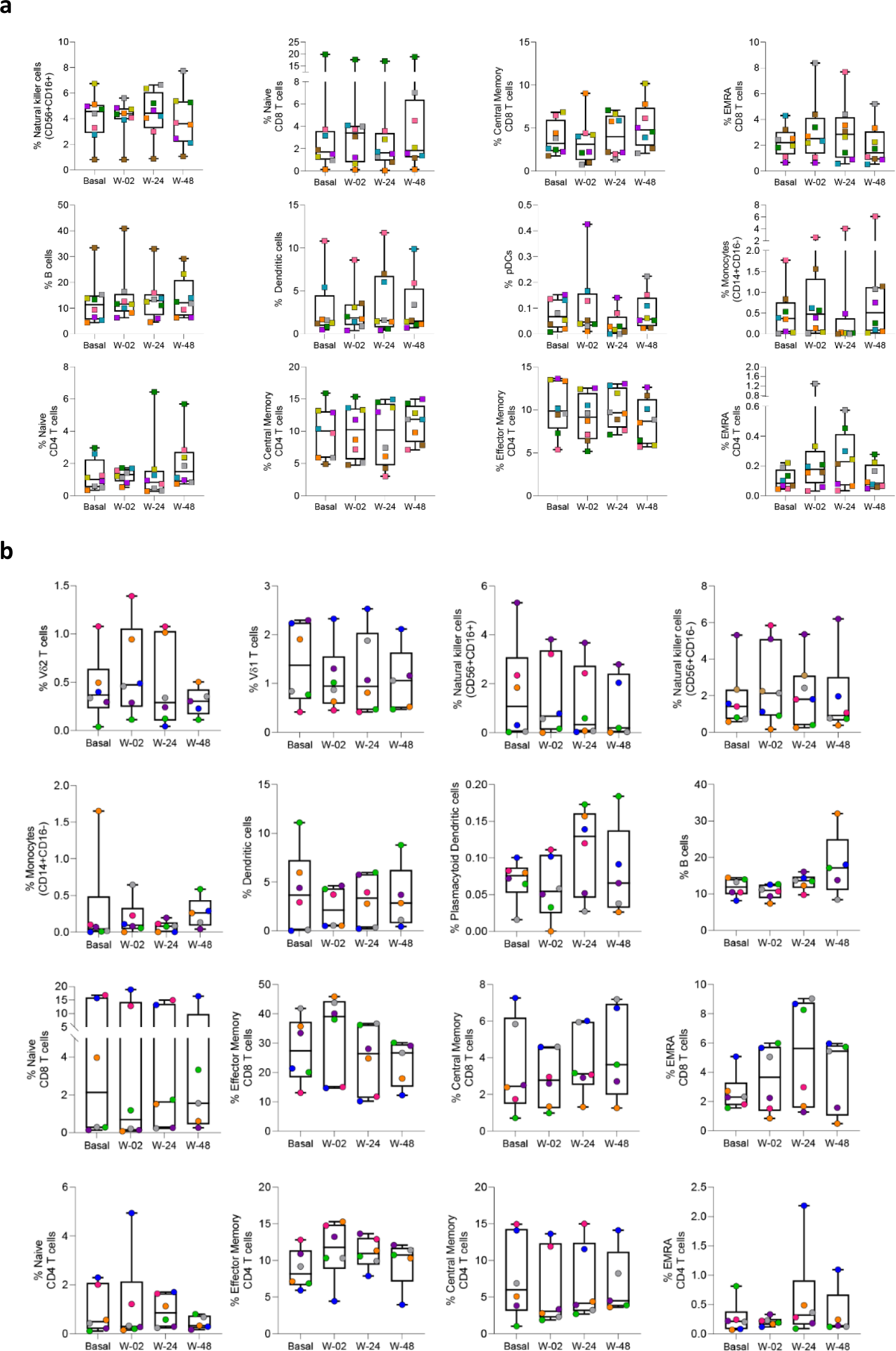
Longitudinal frequencies of circulating cell populations in participants from the A) ALN and B) placebo groups. Wilcoxon signed-rank p-values > 0.05 for all comparisons and adjusted for multiple comparisons using Holm-Bonferroni.

**Figure S8:**
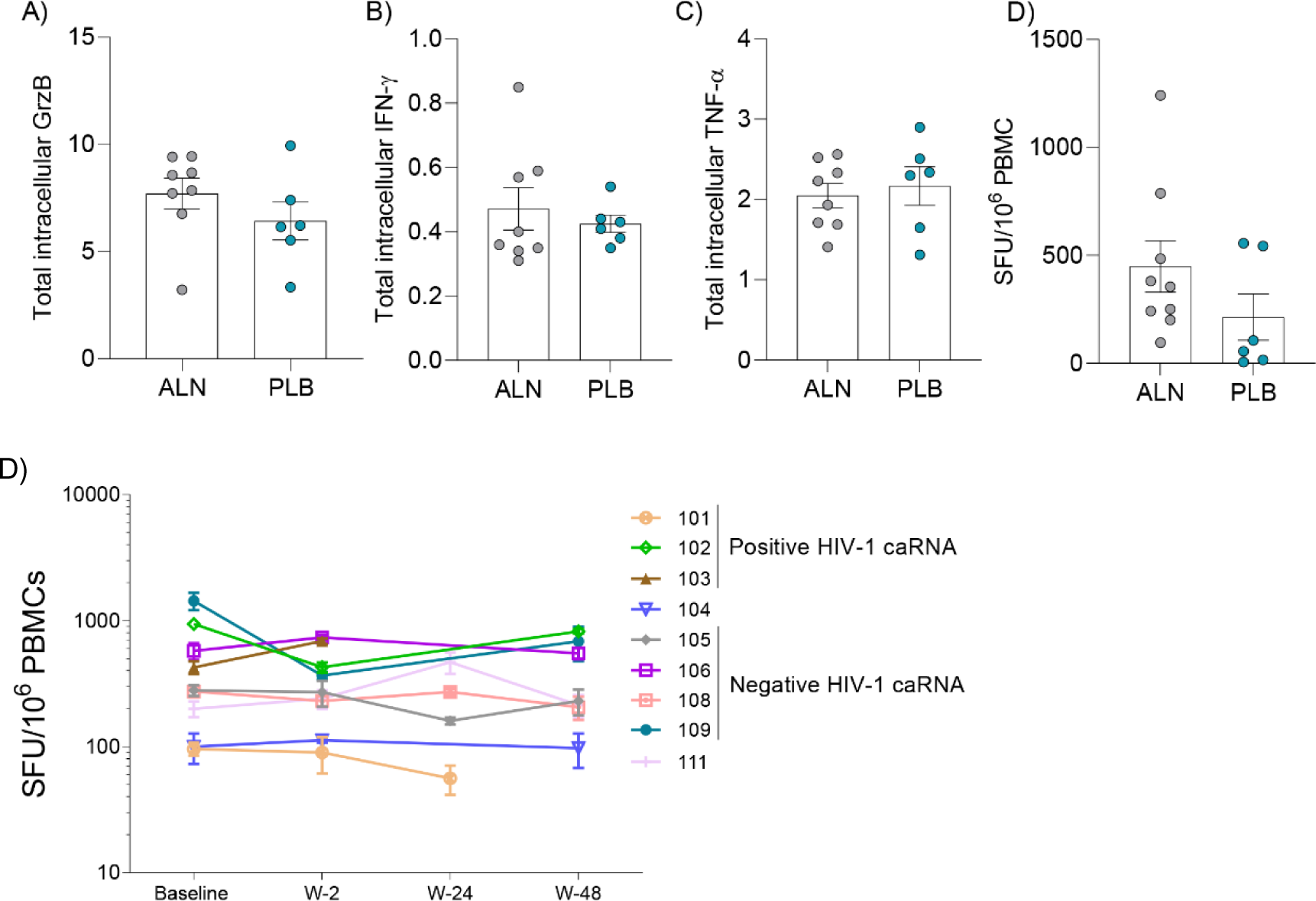
Intracellular functional markers in participants from the ALN and placebo groups. Baseline levels of intracellular production of **A)** granzyme (Grz) B, **B)** IFN-γ, **C)** TNF-α, and **D)** Baseline spot forming units (SFU)/10^6^ PBMCs were comparable between participants who took ALN and PLB (Mann-Whitney U test). **D)** SFU/10^6^ PBMC over time in participants treated with ALN, with legend denoting those with negative versus positive trend in HIV-1 caRNA levels.

**Figure S9:**
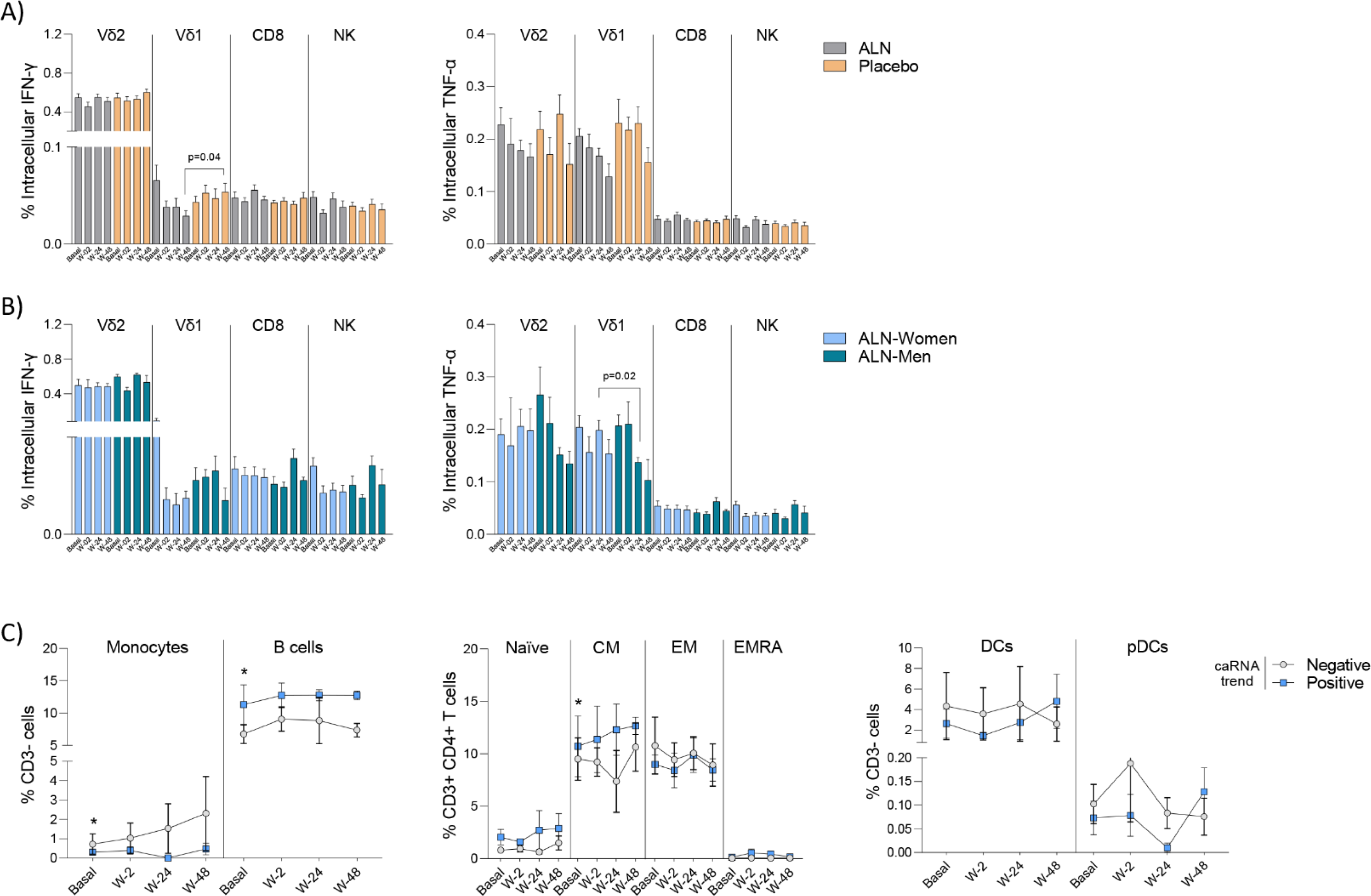
Comparison of IFN-γ and TNF-α production from different effector cell populations (N=8) in A) ALN or placebo groups, B) women and men treated with ALN. C) Comparison of frequencies of circulating cell populations according to the HIV caRNA slope (N=3 negative trend, grey, and N=3 positive trend, blue). Mean ±SEM is presented. Mann-Whitney U test.

